# Biological sex affects human islet gene expression and mitochondrial function in type 2 diabetes

**DOI:** 10.1101/2025.11.10.687716

**Authors:** Sing-Young Chen, Haoning Howard Cen, Charlotte F. Chao, Andrew R. Pepper, James D. Johnson, Elizabeth J. Rideout

## Abstract

The clinical characteristics of type 2 diabetes (T2D) differ between the sexes. For example, the risk of T2D is higher in males than in premenopausal females, whereas the risk of T2D-associated cardiovascular disease is higher in females. However, the sex-dependent mechanisms of T2D pathogenesis remain incompletely understood. Publicly available human islet datasets, such as HPAP and Humanislets.com, offer a valuable tool for uncovering the impact of biological sex on islet structure, gene expression, and function at a scale that was not previously possible. We performed an integrated analysis of data from publicly available sources to identify sex differences in baseline islet characteristics in donors without diabetes and subsequently examined these features in donors who lived with T2D. Among donors without diabetes, female islets had a greater proportion of alpha-cells compared with male islets and showed enriched expression of ribosomal and mitochondrial pathways in both beta- and alpha-cells. Measurements of mitochondrial function in female islets revealed lower spare respiratory capacity compared to male islets. Male and female islets had distinct changes in gene and protein expression in the context of T2D with female islets having greater preservation of insulin content and fewer defects in islet function. Together, these data show female islets have fewer islet impairments in T2D. This highlights the need for detailed mechanistic studies in both sexes to support effective and sex-informed interventions for T2D.

## Introduction

Biological sex influences type 2 diabetes (T2D) incidence, complications, and treatment efficacy. There is a higher diabetes risk in men until approximately 50-60 years of age ^1–4^. This male bias has been observed across all sociodemographic indices ^5^, though we note a female bias in some populations ^6–8^. Relative protection against diabetes in pre-menopausal women has been attributed in part to the protective effects of estrogen ^9,10^. Despite the lower incidence of T2D in premenopausal women, women who live with diabetes are at a higher risk of most T2D complications (e.g. cardiovascular and renal) compared to men ^11–18^. Men and women with T2D also show differential responses to commonly used medications: male sex predicts better HbA1c-lowering efficacy of sulfonylureas, which stimulate beta-cell insulin secretion ^19,20^, and SGLT2 inhibitors, which decrease renal glucose reabsorption ^20,21^, whereas insulin-sensitizing thiazolidinediones and GLP-1 receptor agonists (GLP-1RAs) have greater HbA1c-lowering effects in females ^20–24^.

Sex differences in clinical observations are likely due to a combination of differences in insulin sensitivity and pancreatic beta-cell function. Greater insulin sensitivity in females is well-established and conserved from humans ^25^ to mice ^26^ to fruit flies ^27^. Recent studies suggest that sex differences in pancreatic islets may also play a role. In mice, female islets show greater resilience to endoplasmic reticulum stress than male islets and have higher expression of genes involved in protein synthesis ^28,29^. Sex differences in electrical activity have also been observed in mouse islets: glucose stimulation in female beta-cells results in lower potassium currents leading to a more depolarized membrane potential, more frequent burst-type action potentials, and smaller glucose-induced transient calcium increases ^30,31^. HFD feeding in mice also causes sex-dependent transcriptomic changes ^28^, and female mice are better able to adapt to this metabolic stress through alpha-to-beta cell communication to maintain beta-cell calcium dynamics ^32^. In humans, as in mice, female islets show enriched expression of pathways related to protein synthesis ^29,33^. Beta-cell gene expression is closely linked to epigenetic modifications, which differ between sexes on both sex chromosomes and autosomes, including at T2D candidate genes ^34,35^. *Ex vivo* insulin secretion studies have also suggested that female islets may secrete more insulin than male islets in response to high glucose ^29,33,34^. T2D causes profound changes in gene expression among islets from both sexes, and these changes are also partly sex-dependent ^33,35,36^. However, our understanding of the molecular and functional changes that occur with T2D in female and male islets remains far from complete.

The Human Pancreas Analysis Program (HPAP) consolidates data from donor human islets across multiple institutes. The data collected through this program are uploaded to the publicly available PancDB database, and include measurements of human islet calcium imaging, mass cytometry, islet perifusion, oxygen consumption, and scRNA-seq, among others ^37,38^. Humanislets.com is a repository with functional and omics data on a largely distinct pool of donors, including a wider range of perifusion experiments and a large human islet proteomics dataset ^39^. Importantly, the datasets have enough donors to test for sex differences in many islet and islet cell attributes.

Our analysis of HPAP and Humanislets.com revealed pronounced sex differences in islet composition, RNA and protein levels, and mitochondrial function in islets from donors without diabetes. We further find that gene and protein expression are differentially altered in T2D depending on biological sex, and that male islets exhibited mitochondrial defects that were absent in female islets among donors with T2D. Together, these data demonstrate clear differences in diverse islet characteristics between male and female islets among individuals without T2D, and show sex-dependent changes to islet biology in individuals who lived with T2D.

## Materials and Methods

### Data sources

Publicly available data were downloaded from the Human Pancreas Analysis Program (HPAP) Database, consortia under Human Islet Research Network (RRID:SCR_014393, https://hpap.pmacs.upenn.edu/) ^37,38^ (NIH grant numbers UC4-DK112217 and UC4-DK112232, RRID:SCR_016202), and Humanislets.com ^39^, an initiative of the Alberta Diabetes Institute IsletCore funded by the Canadian Institutes of Health Research, Breakthrough T1D Canada, and Diabetes Canada (5-SRA-2021-1149-S-B/TG 179092). Specific datasets used are described in detail below. To study sex differences within the healthy islet state, we specifically focused on donors without diabetes and aged 15-39 years. This age bracket was chosen to determine sex differences prior to the onset of menopause ^40^ and was used to compare male and female control (non-diabetic) donors only. However, when comparing across control and T2D for males and females, donors of all ages were included. For age-matched data, donors with T2D were matched to donors without diabetes with the closest possible age. Donors with type 1 diabetes were not included in our analyses due to insufficient donor numbers to support sex-specific analysis. Histograms displaying key metadata variables in each dataset are shown in Figure S1^41^.

### Cell type composition

Islet endocrine cell type composition data were available from both HPAP and Humanislets.com. CyToF data ^42^ estimating cell counts for each cell type were obtained from donor metadata available on HPAP, from which endocrine cell type proportions were calculated as described in the CyTOF Staining Workflow and Protocol on HPAP (https://hpap.pmacs.upenn.edu/explore/workflow/imaging-mass-cytometry?protocol=1, protocol accessed 23 September 2025, last updated 11 January 2021). Donors with a total endocrine cell count less than 30-fold lower than the geometric mean of all donors were excluded, as in ^43^. On Humanislets.com, endocrine cell type proportions were estimated based on deconvolution of whole islet proteomics data ^39^ and were downloaded directly.

### Gene expression analysis

Gene expression data from HPAP and Humanislets.com were analyzed to compare the sexes. Transcriptomic data from HPAP included scRNAseq data. To analyze scRNAseq data, whole islet, beta-cell, and alpha-cell expression were first pseudobulked to generate one gene expression profile per donor per cell type using the *Seurat* package ^44^. Pseudobulking overcomes the confounding effects of uneven cell numbers per donor and appropriately treats individual donors, rather than cells, as independent observations. For Humanislets.com, only bulk islet RNAseq was analyzed by sex due to low n’s available for scRNAseq data. Bulk islet proteomics data ^45^ were also downloaded and analyzed from Humanislets.com. Three comparisons were performed: female vs male among donors without diabetes aged 15-39, female control (without diabetes) vs female T2D across all ages, and male control vs male T2D across all ages. For all differential expression analyses, age was included as a covariate.

For whole islet transcriptomics, HPAP pseudobulk data and Humanislets.com bulk RNAseq data were combined and analyzed using *limma* ^46^, with age and dataset as covariates. For HPAP pseudobulk beta-cell, HPAP pseudobulk alpha-cell, and Humanislets.com proteomics data, *limma* ^46^ was used with age as a covariate. Gene Set Enrichment Analysis (GSEA) was performed on differential expression analysis output to identify key pathways using the clusterProfiler package ^47^, in which the direction signed -log10 *p*-values were used as the rank scores of the genes. To improve pathway visualization, pathways were considered redundant if their core enrichment genes exceeded a Jaccard index of 0.6. When selecting pathways for visualization, redundant pathways were first refined by omitting the pathway with fewer gene components. For visualization of pathways, we grouped redundant or similar pathways together by using Leiden clustering^48^ with Jaccard index as similarity measurement between pathways. For comparisons between female and male control donors aged 15-39, pathways with adjusted *p*-value < 0.05 were included in visualization. For comparisons between control and T2D donors, pathways with adjusted *p*-value < 0.0001 were included in visualization, but all pathways with adjusted *p*-value < 0.05 were included in tables.

### Insulin content

Islet insulin content data were downloaded from perifusion datasets on HPAP. There were two sets of islet perifusion data with corresponding insulin content data. The data from the University of Pennsylvania (UPenn) were presented per islet using a lysis buffer containing EDTA and detergent ^49,50^. The data from Vanderbilt University were presented per islet equivalent (IEQ), using acid-ethanol extraction to lyse cells and isolate insulin. Acid-ethanol extraction is widely used in the field to extract insulin from biological samples at maximum yields ^51–55^, leveraging the strong acid to avoid the insulin precipitation pH range of ∼4.5-6.5 ^52,56^ and ethanol to minimize the contaminating influences of exocrine impurities ^57,58^ while retaining insulin in solution ^59^. Humanislets.com also used acid-ethanol extraction and presented insulin content data per IEQ. These data were obtained from the provided metadata. Given the consistent methods and units, the Vanderbilt and Humanislets.com data were pooled for combined analyses with dataset as an additional covariate.

### Oxygen consumption

Islet oxygen consumption in response to stimuli was available from both HPAP and Humanislets.com. On Humanislets.com, oxygen consumption data were obtained using a Seahorse Bioanalyzer (Agilent) and calculated parameters were directly downloaded (spare respiratory capacity = maximum uncoupled respiration – baseline, maximum respiratory capacity = maximum uncoupled respiration – minimum non-mitochondrial respiration, response to high glucose = mean respiration in 16.7 mM glucose – mean respiration in 3 mM glucose, baseline = last basal respiration) ^39^. As three readings per donor were available, outliers were identified using Grubb’s test and the average per donor was calculated. On HPAP, these data were obtained using a chemical oxygen probe (Oxyphor G3). Raw data were downloaded and parameters were calculated using the same formulae as used on Humanislets.com. The HPAP and Humanislets.com data were combined to assess potential sex differences among donors aged 15-39 without diabetes. Due to the low number of T2D donors in the Humanislets.com dataset, only the HPAP dataset was used for assessing the impact of T2D status.

### Intracellular calcium

Intracellular calcium imaging data using the ratiometric dye Fura-2 were available only from HPAP. Raw data were downloaded and processed as follows: 1) only regions that responded to high glucose were included (response counted if 16.7 mM glucose stimulated an increase in baseline-corrected calcium signal of at least 50%); 2) after filtering for glucose response, signals from multiple regions for each run were averaged; 3) runs without a specified time of first stimulus addition or without high glucose were excluded; 4) to account for different time periods between stimuli, only the first x minutes after stimulus addition were counted for each run, where x is the minimum period of time between that stimulus and the next, for any of the runs; 5) responses were averaged for each individual donor and summary statistics were calculated.

### Dynamic hormone secretion

HPAP provides dynamic hormone secretion data from islet perifusion experiments performed at two sites: UPenn and Vanderbilt. For the UPenn data, areas under the curve (AUC) were calculated for the periods of perifusion during which islets were exposed to each stimulus, for both insulin and glucagon secretion. For the Vanderbilt data, there was a visible delay between stimulus addition and response. Therefore, the AUC calculated for each stimulus was determined based on the time course tracings: 12-39 min for 16.7 mM glucose, 63-87 min for 16.7 mM glucose + IBMX, 93-111 min for 1.7 mM glucose + epinephrine, and 123-138 min for KCl. Humanislets.com contains islet perifusion data measuring insulin secretion in response to glucose, leucine, and a mix of the free fatty acids oleate and palmitate ^45^. For each Humanislets.com perifusion experiment, AUC were calculated for the periods of perifusion during which islets were exposed to each stimulus. As the HPAP UPenn and Humanislets.com datasets used equivalent units and similar glucose concentrations (3 mM for low glucose), these were combined for analyses and summary statistics were taken, including mean insulin secretion at 3 mM glucose, peak insulin secretion at high glucose (15 mM for Humanislets.com, 16.7 mM for HPAP UPenn), peak insulin secretion at 30 mM KCl, and stimulation index (ratio of peak secretion at high glucose over mean secretion at 3 mM glucose).

### Statistics

Unless otherwise specified, to determine statistical differences due to sex and disease, ANCOVA testing was performed with age as covariate. When multiple datasets were combined, dataset was also included as a covariate. Multiple comparisons were performed using Tukey’s correction and the *emmeans* R package. For age-matched data, donors with T2D were matched to donors without diabetes with the closest possible age within each sex using the *MatchIt* package, and statistical comparisons were performed to assess the effect of T2D status only. As some significant age differences remained in some cases, age was still included as a covariate for statistical analyses of age-matched data. For comparing distributions in Figure S1, the two-sample Kolgomorov-Smirnov test was used. Unless otherwise mentioned, a *p*-value threshold of 0.05 (or adjusted *p*-valued threshold where relevant) was used. All relevant scripts for data processing and graphing are available at https://github.com/singyoungchen/sex-differences-human-islet-characteristics-.

## Results

### Female islets contain a greater proportion of alpha-cells and a smaller proportion of beta-cells

We first examined islet cell type composition in male and female donors without T2D aged 15-39 years. Female islets had a significantly higher alpha-cell proportion compared to male islets in both the HPAP and Humanislets.com datasets (Figure 1A-B). Consistent with increased alpha-cell proportion, beta-cell proportion was lower in female islets compared with male islets, a difference that was statistically significant in the Humanislets.com data and which showed a non-significant but similar trend (*p*=0.11) in the HPAP data (Figure 1C-D). The proportion of other endocrine cells was significantly higher in males than females in HPAP (Figure 1E) but not Humanislets.com (Figure 1F). When donors of all ages were considered together, however, the trends we observed were not consistent between the HPAP CyToF results and the Humanislets.com cell type proportion estimates. The HPAP data suggested that, among donors without diabetes of all ages, female islets showed higher alpha-cell (*p*<0.05, Figure 1G), lower beta-cell (*p*=0.072, Figure 1H), and lower non-alpha, non-beta cell (*p*=0.083, Figure 1I) proportions, consistent with observations from donors in the 15-39 age bracket. In contrast, when including data for donors of all ages from Humanislets.com, we observed no sex difference in endocrine cell proportions between female and male donors without T2D (Figure 1J-L).

**Figure 1.**
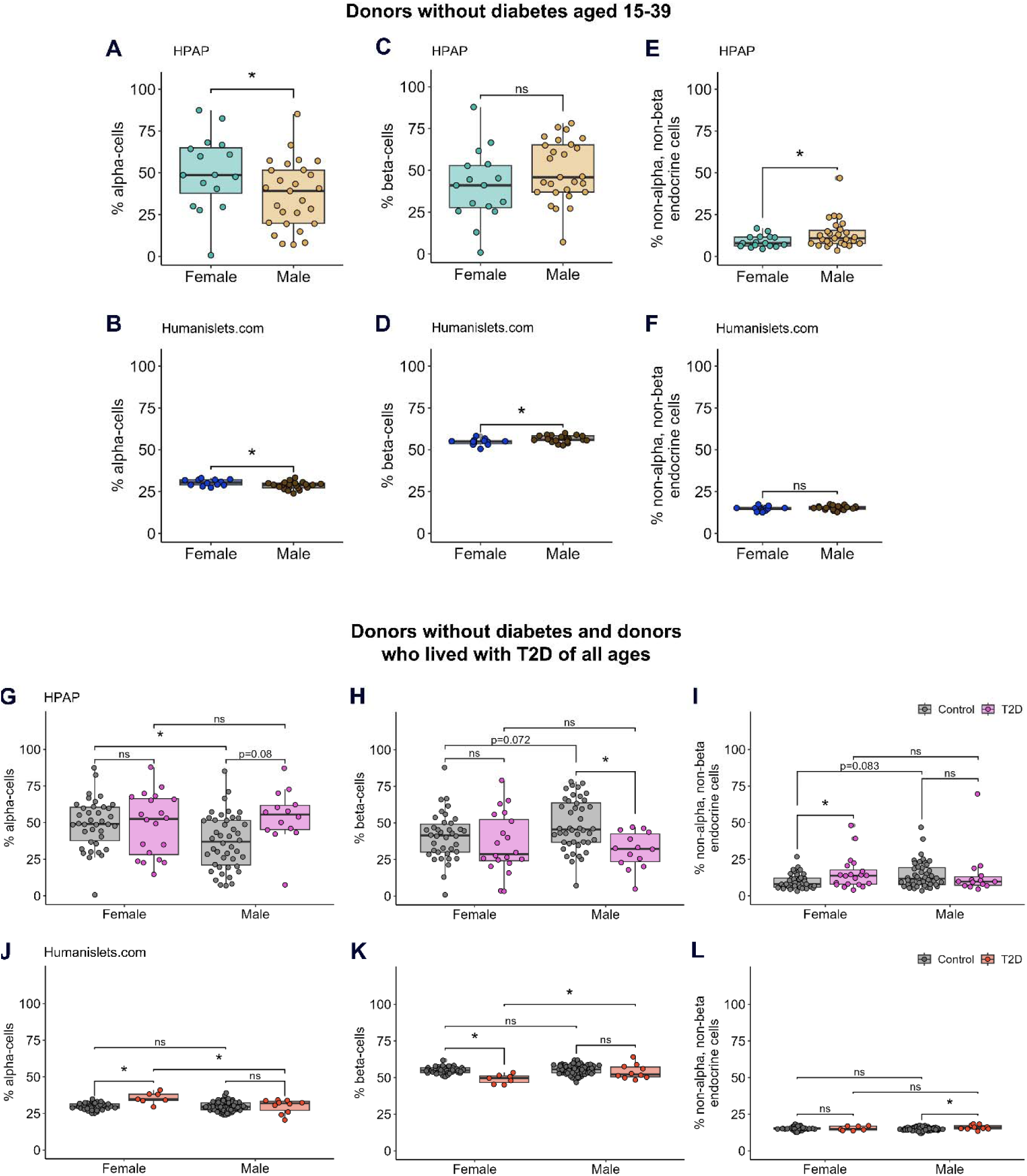
Islet endocrine cell proportions. Proportions were determined by CyToF in the HPAP dataset or deconvolution from whole islet proteomics in the Humanislets.com dataset. For islets from donors without diabetes aged 15-39, alpha-cell proportions from HPAP (A) and Humanislets.com (B), beta-cell proportions from HPAP (C) and Humanislets.com (D), and non-alpha, non-beta endocrine cell proportions from HPAP (E) and Humanislets.com (F). For islets from donors without diabetes and donors with T2D of all ages, alpha-cell (G), beta-cell (H), and non-alpha, non-beta endocrine cell (I) proportions from HPAP and alpha-cell (J), beta-cell (K), and non-alpha, non-beta endocrine cell (L) from Humanislets.com. * indicates *p*<0.05, ns = not significant, *p*_≥_0.05.

We next considered endocrine cell proportions in donors who lived with T2D. The HPAP data showed that T2D was associated with a trend toward increased alpha-cell proportion (*p*=0.08) and a significant decrease in beta-cell proportion (*p*<0.05) in males but not females (Figure 1G-H). The HPAP data also showed a T2D-associated increase in the proportion of other endocrine cells in females but not males (Figure 1I). In contrast, the Humanislets.com data showed an increase in alpha-cell proportion and a decrease in beta-cell proportion in female, but not male, donors who lived with T2D (Figure 1J-K). In the Humanislets.com dataset, the proportion of other endocrine cells was significantly increased with T2D in males only (Figure 1L). These trends remained in age-matched data (Figure S2^41^). While the reason for this discrepancy is unclear, it may reflect the different techniques used by each team to obtain cell proportion estimates (antibody-based identification in HPAP, estimation based on marker protein expression data in Humanislets.com). Thus, while alpha-cell proportion was elevated among females in young donors without diabetes, we observed no consistent sex difference in donors who lived with T2D across datasets.

### Islets from young female donors show higher expression of protein synthesis genes

We next checked for sex differences in baseline gene expression among islets from donors aged 15-39 without diabetes by combining pseudobulk scRNAseq data from HPAP with bulk islet RNAseq from Humanislets.com. After conducting differential expression analysis to compare female and male donors (Table 1, S1^41^), we performed GSEA. We found 23 female-biased pathways and 16 male-biased pathways (Figure 2A, Table S2^41^). Pathways related to ribosomal biogenesis and protein synthesis showed a strong female bias (Figure 2A, Table S2^41^). The potential for enhanced protein synthesis capacity in female islets has previously been shown by us ^29^, and others ^33^, and is likely associated with the effects of estrogen in alleviating ER stress and preserving protein synthesis ^60–63^. Mitochondrial pathways were also female-enriched (Figure 2A, Table S2^41^). Male-biased pathways included those related to cell division and histone modification (Figure 2A, Table S2^41^). The HPAP scRNAseq data also allowed us to perform sex-based analysis of beta-cell- and alpha-cell-specific pseudobulk data. Both cell types showed a female-biased enrichment of ribosomal and mitochondrial gene pathways and a male-biased enrichment of mitosis and histone modification pathways (Figure S3^41^, Table S2^41^).

**Figure 2.**
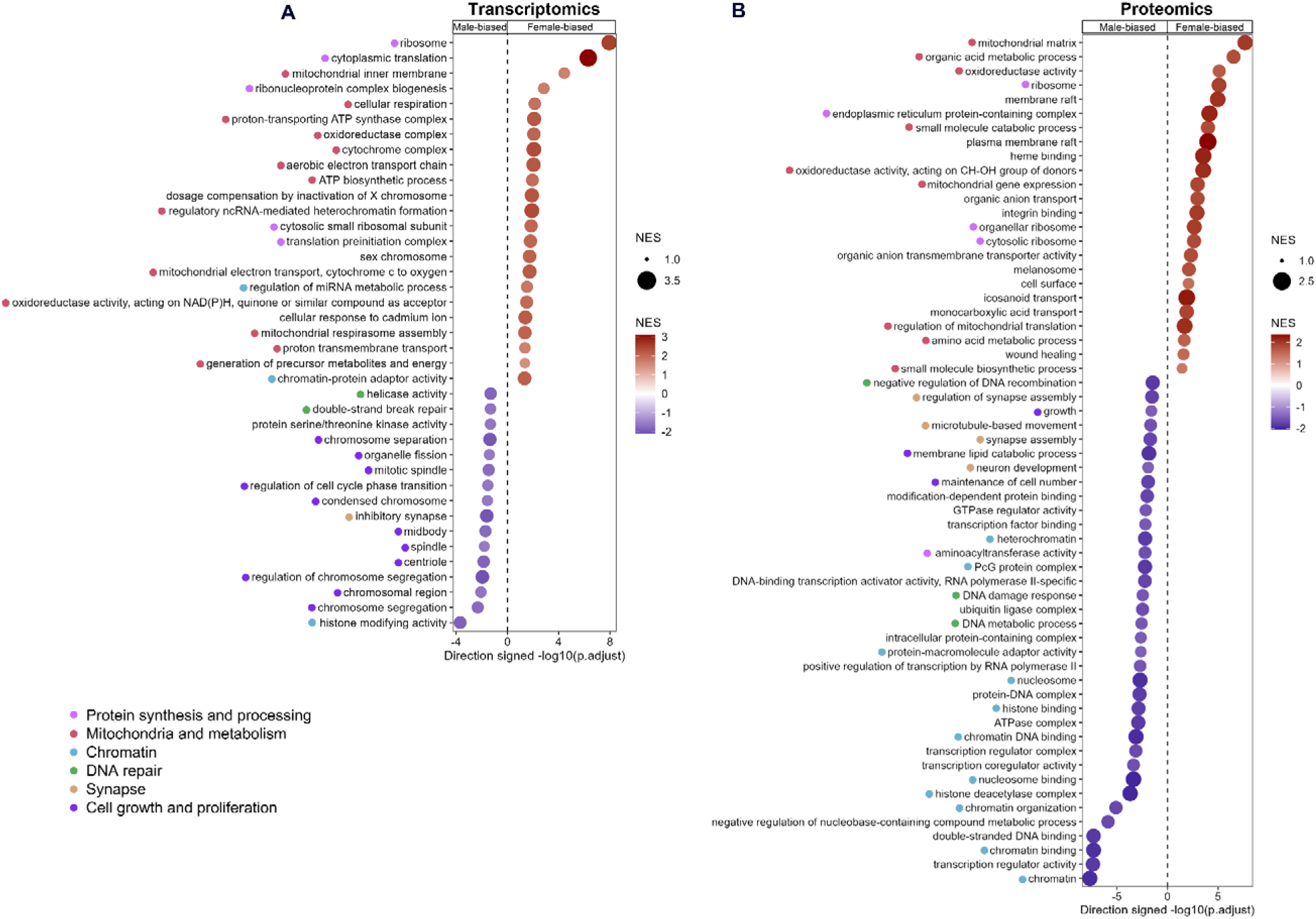
Gene and protein expression analysis of islets from female versus male donors without diabetes aged 15-39. Significantly altered pathways according to sex in GSEA analysis among combined transcriptomics datasets (HPAP pseudobulk scRNAseq and Humanislets.com bulk islet RNAseq) (A). Top 60 GO pathways that were significantly altered by sex in GSEA analysis in Humanislets.com bulk islet proteomics (B). n = 7 female, 17 male for HPAP scRNAseq, 8 female, 18 male for Humanislets.com bulk RNAseq, and 12 female, 21 male for Humanislets.com proteomics. NES = normalized enrichment score. Pink dots indicate pathways associated with protein synthesis and processing, red dots indicate pathways associated with mitochondrial regulation and function, blue dots indicate pathways associated with chromatin and chromatin remodeling, green dots indicate pathways associated with DNA repair, yellow dots indicate pathways associated with synapses and synapse formation, and purple dots indicate pathways associated with cell growth and proliferation.

**Table 1.**
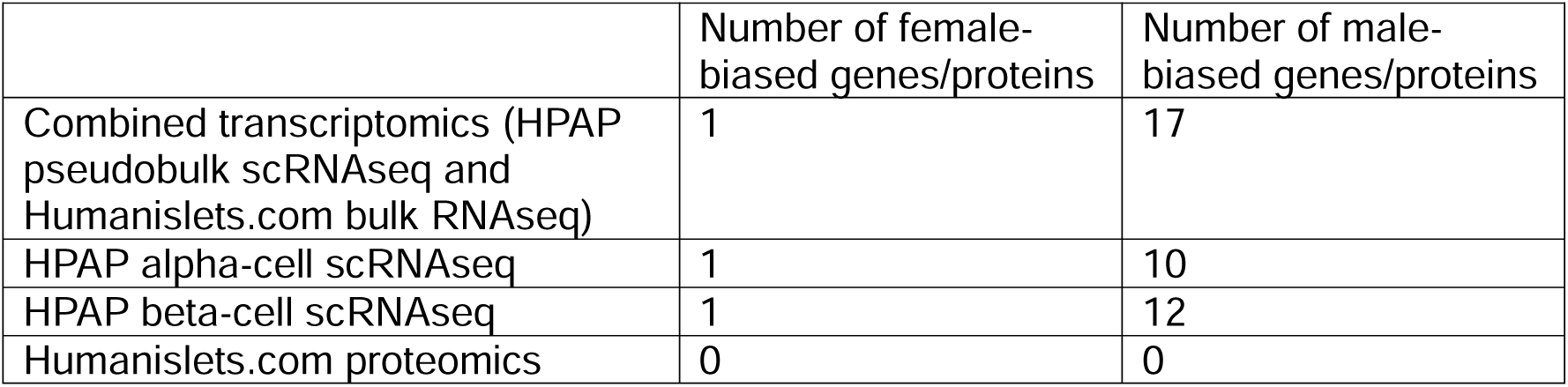
Total genes or proteins significantly altered according to biological sex in differential expression analysis among islets from donors without diabetes aged 15-39.

Humanislets.com includes, to our knowledge, the largest open-source whole islet proteomics dataset currently available. We conducted differential expression analysis to compare protein levels in female and male donors without diabetes aged 15-39 (Table S3^41^) followed by GSEA. We found female-biased abundance of proteins related to ribosomes and mitochondria, as well as lipid transport and the plasma membrane (Figure 2B, Table S4^41^). Male-biased pathways included those related to chromatin modification and transcriptional regulation (Figure 2B, Table S4^41^). Female-biased and male-biased pathways identified in proteomics included many similar pathways as those identified in transcriptomics (Table 2). Thus, the female bias in expression of proteins related to protein synthesis and mitochondrial function was highly consistent between our RNAseq and proteomics data.

**Table 2.** Categories of female-biased and male-biased GO pathways identified by GSEA in islets from donors without diabetes aged 15-39. Values indicate percentage of significant pathways in that category (significant pathways in category/total significant pathways).

|  | **Combined transcriptomics** | **Proteomics** |
| --- | --- | --- |
| *Female-biased pathways* |  |  |
| Mitochondria | 47.8% (11/23) | 13.2% (12/91) |
| Ribosome and protein synthesis | 21.7% (5/23) | 15.4% (14/91) |
| Metabolic process | 8.7% (2/23) | 19.8% (18/91) |
| Oxidative stress response | 4.4% (1/23) | 4.4% (4/91) |
| Metabolite transport | 0 | 23.1% (21/91) |
| Plasma membrane | 0 | 13.2% (12/91) |
| Sex chromosome | 8.7% (2/23) | 0 |
| Wound healing | 0 | 2.2% (2/91) |
| Endoplasmic reticulum and Golgi | 0 | 6.6% (6/91) |
| Chromatin | 8.7% (2/23) | 0 |
| Melanosome | 0 | 2.2% (2/91) |
| *Male-biased pathways* |  |  |
| Chromatin | 6.3% (1/16) | 21.0% (25/119) |
| Cell growth and proliferation | 68.9% (11/16) | 6.7% (8/119) |
| DNA repair | 12.5% (2/16) | 9.2% (11/119) |
| Synapse | 6.3% (1/16) | 18.5% (22/119) |
| Cell regulation | 6.3% (1/16) | 0.8% (1/119) |
| Transcriptional regulation | 0 | 28.6% (34/119) |
| Protein synthesis, modification, and degradation | 0 | 9.2% (11/119) |
| Lysosome | 0 | 0.8% (1/119) |
| ATPase and GTPase | 0 | 3.4% (4/119) |

### T2D is associated with differential shifts in gene and protein expression depending on sex

We next compared islet gene expression between donors with and without diabetes to evaluate any sex bias in gene expression changes associated with T2D. In the combined transcriptomics dataset, 3 genes were significantly altered with T2D in females and 431 in males (Table 3, S5-6^41^). GSEA identified more pathways significantly altered with T2D in males than in females (Table 4, S7-8^41^). Among the pathways increased with T2D in female islets were pathways related to transcriptional regulation and histone modification (Figure 3A, Table S7^41^). In male islets, pathways related to ribosomes and inflammation were increased with T2D (Figure 3B, Table S8^41^). Pathways decreased with T2D in female islets were dominated by pathways such as ribosomal and mitochondrial gene pathways (Figure 4A, Table S7^41^). In males, T2D was associated with a decrease in pathways related to mitochondria, vesicle processing and secretion, and microtubules (Figure 4B, Table S8^41^). Our data therefore suggest that there were distinct T2D-associated changes in gene expression between the sexes in whole islets.

**Figure 3.**
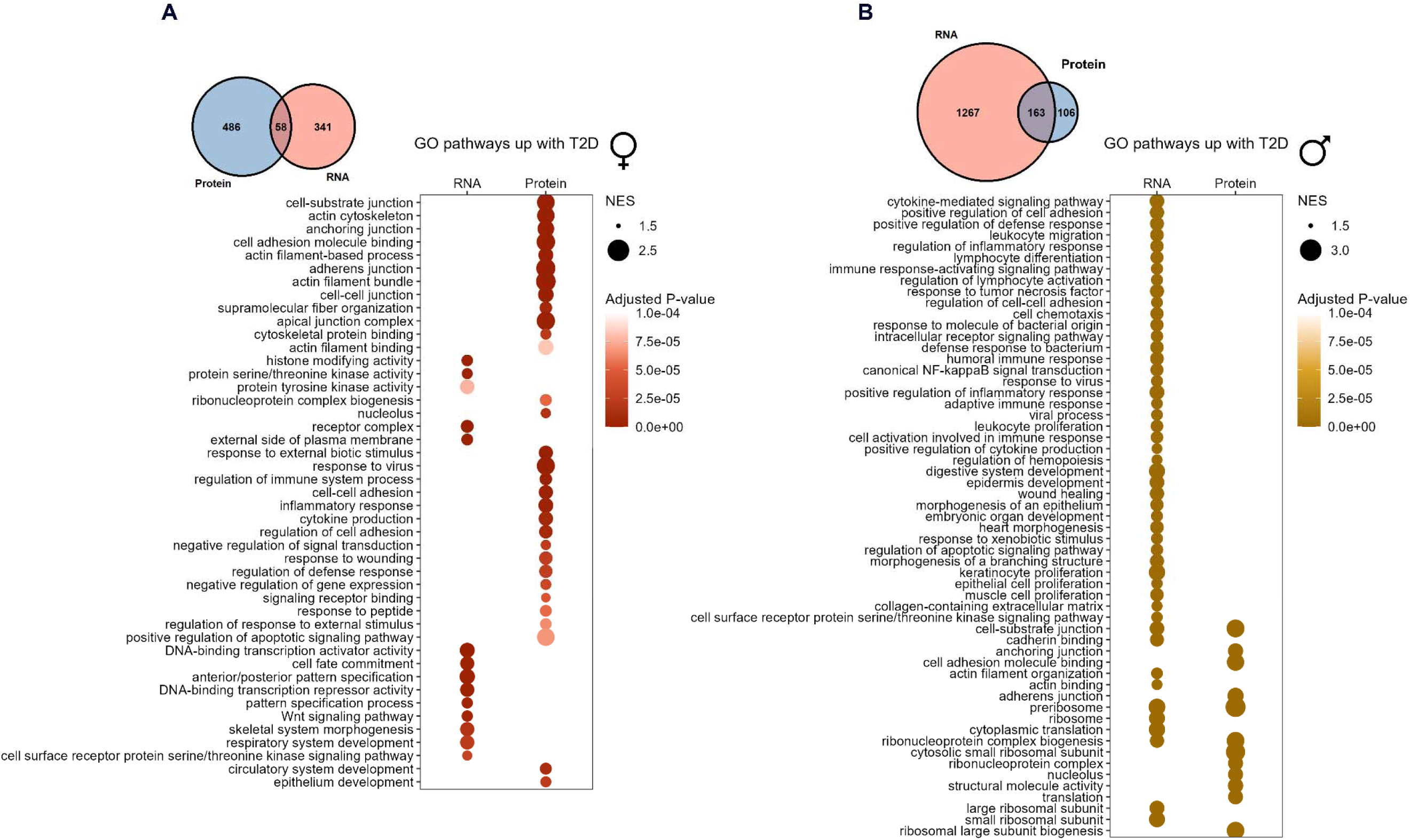
GO pathways upregulated with T2D in islets from female and male donors of all ages. Pathways that were significantly increased with T2D in female islets in GSEA analysis at *p*<0.0001 among combined transcriptomics datasets (HPAP pseudobulk scRNAseq and Humanislets.com bulk islet RNAseq) and Humanislets.com proteomics (A). Top 60 pathways that were significantly increased with T2D in male islets in GSEA analysis at *p*<0.0001 among combined transcriptomics datasets and Humanislets.com proteomics (B). Venn diagrams show common and distinct pathways that were significantly increased with T2D at *p*<0.05 in the combined transcriptomics versus Humanislets.com proteomics datasets. For HPAP scRNAseq, n = 16 female control, 11 female T2D, 24 male control, and 7 male T2D donors. For Humanislets.com bulk RNAseq, n = 36 female control, 5 female T2D, 66 male control, and 10 male T2D donors. For Humanislets.com proteomics, n = 40 female control, 7 female T2D, 77 male control, and 10 male T2D donors. NES = normalized enrichment score.

**Figure 4.**
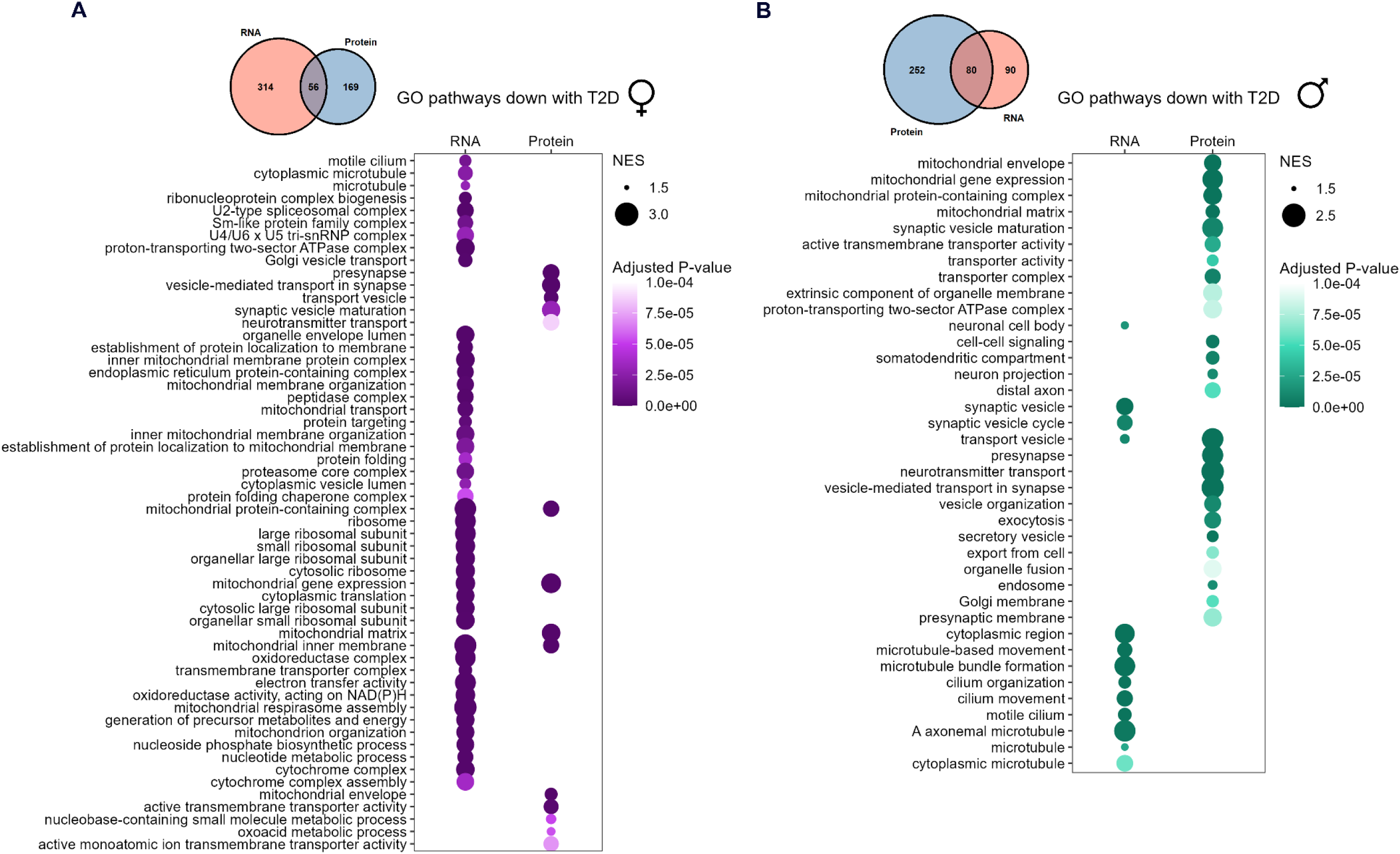
GO pathways downregulated with T2D in islets from female and male donors of all ages. Pathways that were significantly decreased with T2D in female islets in GSEA analysis at *p*<0.0001 among combined transcriptomics datasets (HPAP pseudobulk scRNAseq and Humanislets.com bulk islet RNAseq) and Humanislets.com proteomics (A). Pathways that were significantly decreased with T2D in male islets in GSEA analysis at *p*<0.0001 among combined transcriptomics datasets and Humanislets.com proteomics (B). Venn diagrams show common and distinct pathways that were significantly decreased with T2D at *p*<0.05 in the combined transcriptomics versus Humanislets.com proteomics datasets. For HPAP scRNAseq, n = 16 female control, 11 female T2D, 24 male control, and 7 male T2D donors. For Humanislets.com bulk RNAseq, n = 36 female control, 5 female T2D, 66 male control, and 10 male T2D donors. For Humanislets.com proteomics, n = 40 female control, 7 female T2D, 77 male control, and 10 male T2D donors. NES = normalized enrichment score.

**Table 3.** Total genes or proteins significantly altered according to T2D status in differential expression in islets from donors of all ages.

|  | Number of genes/proteins increased with T2D in females | Number of genes/proteins increased with T2D in males | Number of genes/proteins decreased with T2D in females | Number of genes/proteins decreased with T2D in males |
| --- | --- | --- | --- | --- |
| Combined transcriptomics (HPAP pseudobulk scRNAseq and Humanislets.com bulk RNAseq) | 0 | 310 | 3 | 121 |
| HPAP alpha-cell scRNAseq | 0 | 0 | 0 | 0 |
| HPAP beta-cell scRNAseq | 0 | 6 | 0 | 8 |
| Humanislets.com proteomics | 371 | 299 | 300 | 413 |

**Table 4.**
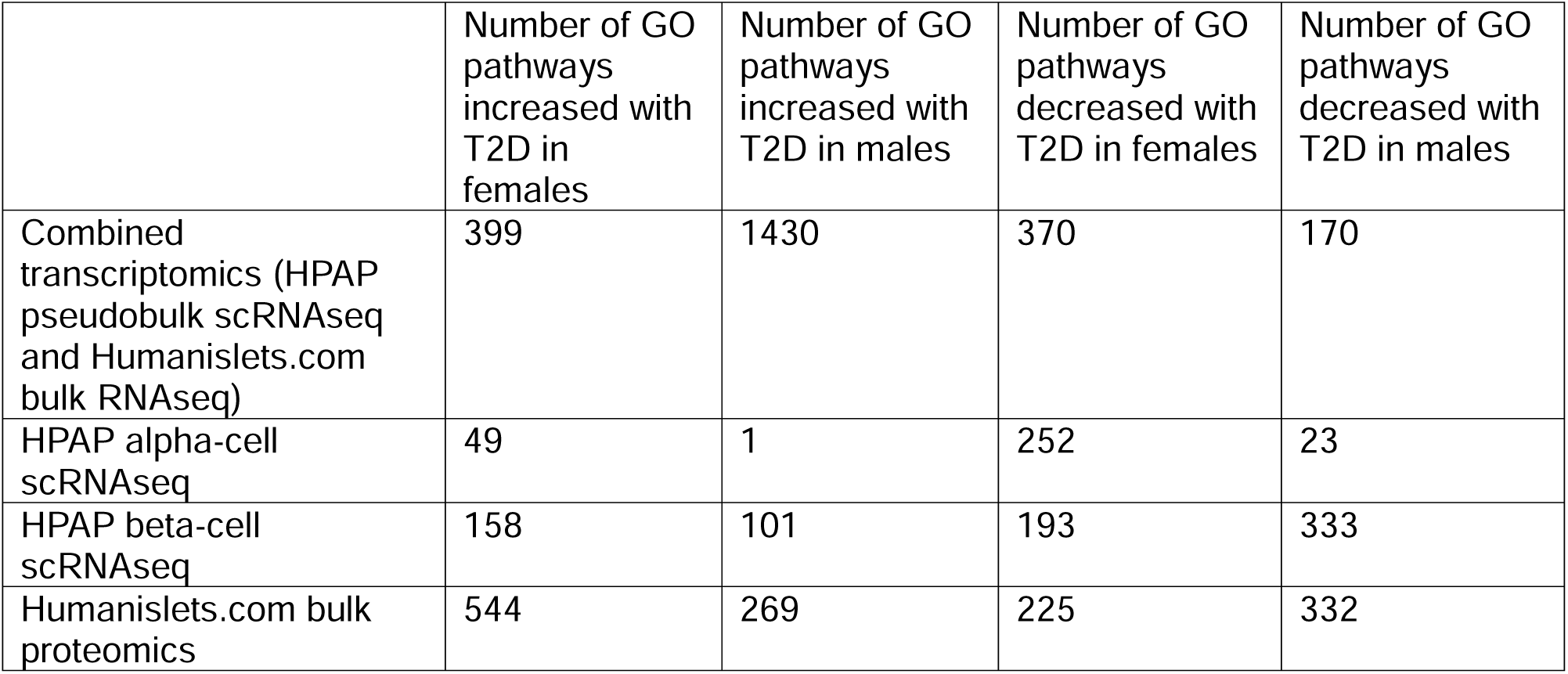
Total GO pathways significantly altered according to T2D status in GSEA in islets from donors of all ages.

|  | Number of GO pathways increased with T2D in females | Number of GO pathways increased with T2D in males | Number of GO pathways decreased with T2D in females | Number of GO pathways decreased with T2D in males |
| --- | --- | --- | --- | --- |
| Combined transcriptomics (HPAP pseudobulk scRNAseq and Humanislets.com bulk RNAseq) | 399 | 1430 | 370 | 170 |
| HPAP alpha-cell scRNAseq | 49 | 1 | 252 | 23 |
| HPAP beta-cell scRNAseq | 158 | 101 | 193 | 333 |
| Humanislets.com bulk proteomics | 544 | 269 | 225 | 332 |

We similarly analyzed beta-cell and alpha-cell pseudobulk scRNAseq data from HPAP. In females, T2D was associated with positive enrichment of transcriptional regulation pathways and negative regulation of mitochondrial and ribosomal pathways in both cell types (Figure S4A-B^41^, Table S7^41^). In males, T2D was associated with an increase in RNA splicing and ribosome pathways, and a decrease in angiogenesis and extracellular matrix pathways in beta-cells (Figure S4C^41^, Table S8^41^). In alpha-cells among males, no significant pathways were detected at the *p* < 0.0001 threshold (Figure S4D^41^), suggesting that T2D may be associated with greater changes in alpha-cell gene expression in female islets than in male islets.

At the protein level, T2D was associated with altered expression of 671 proteins in female islets and 712 proteins in male islets (Table 3, Table S9-10^41^). GSEA showed that T2D was associated with enrichment of pathways related to cell adhesion in both sexes (Figure 3A-B, Table S11-12^41^). In female islets, T2D was also associated with enrichment of pathways related to the inflammatory response and cytoskeleton organization (Figure 3A, Table S11^41^). In male islets, T2D was associated with an enrichment of cytosolic ribosomal pathways (Figure 3B, Table S12^41^), consistent with RNAseq data (Figure 3B, Table S8^41^). Pathways that were decreased with T2D were similar between males and females, and included those related to vesicle trafficking and exocytosis, as well as mitochondrial components (Figure 4A-B, Table S11-12^41^). Unlike in non-T2D islets, pathways identified based on protein expression largely did not overlap with pathways from our gene expression data, highlighting the importance of collecting multiple data types. Taken together, our analysis reveals sex differences in both islet gene and protein expression in donors without diabetes and individuals who lived with T2D.

### Female islets may show greater preservation of insulin content than male islets in T2D

Healthy beta-cells have a remarkable capacity to rapidly increase insulin biosynthesis upon glucose stimulation ^64^. In the context of T2D, islet insulin content has often ^65^, but not always ^45,66^, been reported to decrease. Given the sex differences in ribosome- and protein synthesis-related pathways, we compared islet insulin content in male and female islets.

The Humanislets.com and Vanderbilt datasets both used the acid-ethanol method to obtain insulin content data, including from detergent resistant granules, and normalized to IEQ; therefore, these two datasets were analyzed both together and independently. The UPenn dataset involved a detergent-based method to obtain insulin content, possibly reflecting more recently synthesized insulin and less granule-stored insulin, and normalized per islet; this was analyzed alone. Among donors without diabetes aged 15-39, no significant sex difference in insulin content was observed either in the combined Humanislets.com and Vanderbilt data (Figure 5A), or when these datasets were analyzed independently (Figure S5A-B^41^). The UPenn data showed a trend (*p*=0.068) toward higher insulin content in female islets (Figure 5B). Among donors without diabetes of all ages, Humanislets.com and Vanderbilt data found no sex difference in insulin content (Figure 5C, S5C-D^41^), whereas UPenn insulin content data showed significantly higher islet insulin content in females (Figure 5D). Thus, insulin solubilized using the detergent method showed higher insulin content in female islets among donors without diabetes, but not insulin solubilized by acid-ethanol.

**Figure 5.**
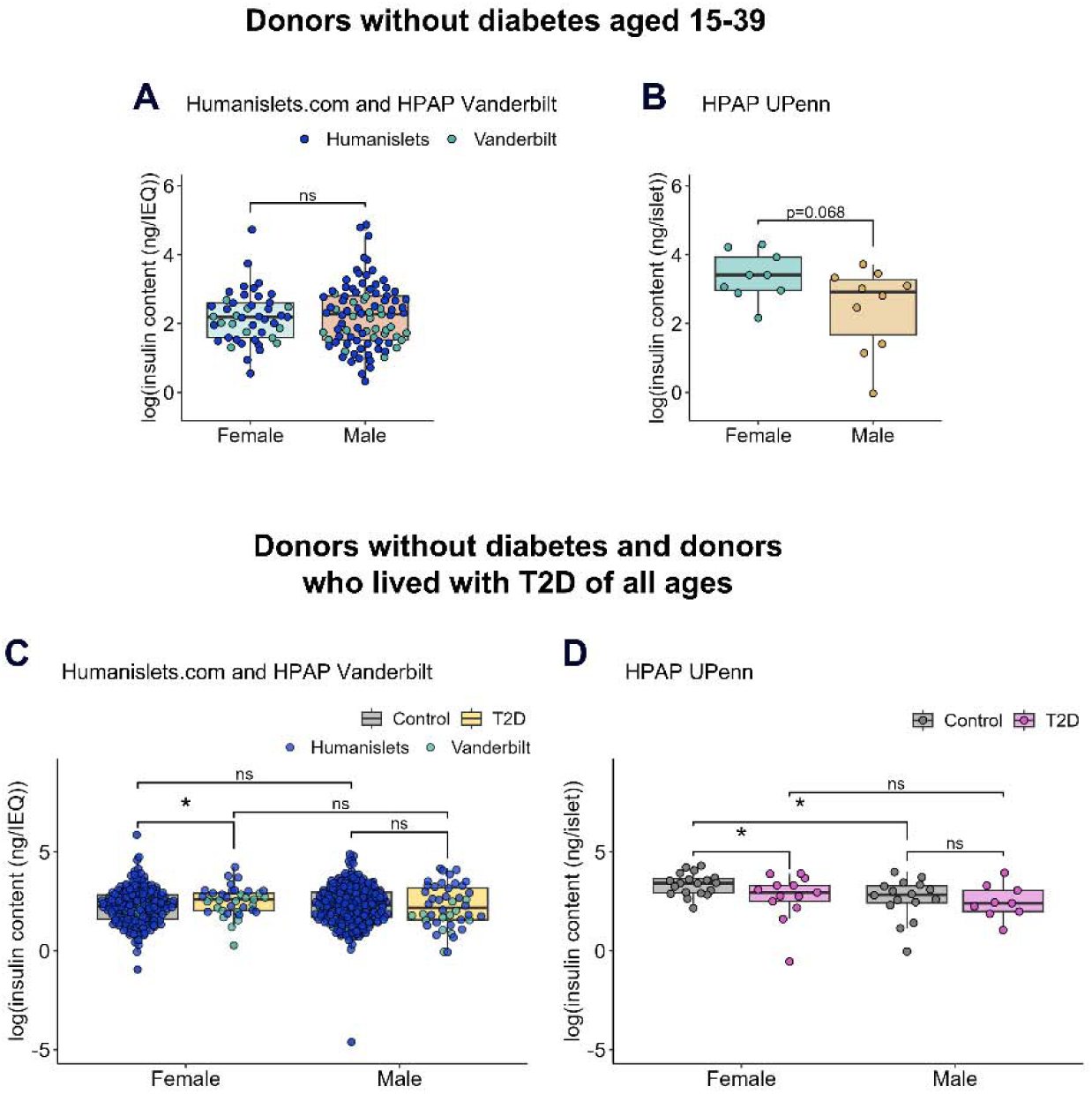
Islet insulin content. Data shown from combined Humanislets.com and HPAP Vanderbilt (A), and HPAP UPenn (B) for donors without diabetes aged 15-39, and from combined Humanislets.com and HPAP Vanderbilt (C) and HPAP UPenn (D) for donors without diabetes and donors with T2D of all ages. * indicates *p*<0.05, ns = not significant, *p*_≥_0.05.

With respect to the effect of T2D on islet insulin content, data from Vanderbilt and Humanislets.com data showed that T2D was associated with increased insulin content in females (Figure 5C, S5C-D^41^), which remained the case when data were age-matched for Vanderbilt, though not Humanislets.com (Figure S6A-F^41^). This finding was not reproduced in the UPenn data, in which T2D was associated with decreased insulin content among female donors (Figure 5D, S6G-H^41^). Male donors who lived with T2D did not have significantly altered islet insulin content compared to male donors without diabetes across all three datasets (Figure 5C-D, S5C-D, S6^41^). Therefore, T2D may be associated with increased acid-ethanol-solubilized insulin but decreased detergent-solubilized insulin specifically in female islets.

### Islets from female donors who lived with T2D have preserved mitochondrial activity

We next examined whether sex differences were present in islet oxygen consumption rate (OCR) data, which reflects mitochondrial function ^67–69^. For donors without diabetes aged 15-39, we combined HPAP and Humanislets.com OCR data (Figure 6A-B) to analyze key parameters related to mitochondrial function. Neither basal nor maximal respiration was different between the sexes (Figure 6C-D). Spare respiratory capacity refers to the difference between maximal respiratory capacity and basal respiration and reflects cells’ ability to ramp up respiration in response to increased energy demand. Female islets had lower spare respiratory capacity than male islets (Figure 6E). Similar trends were observed when datasets were analyzed individually (Figure S7).

**Figure 6.**
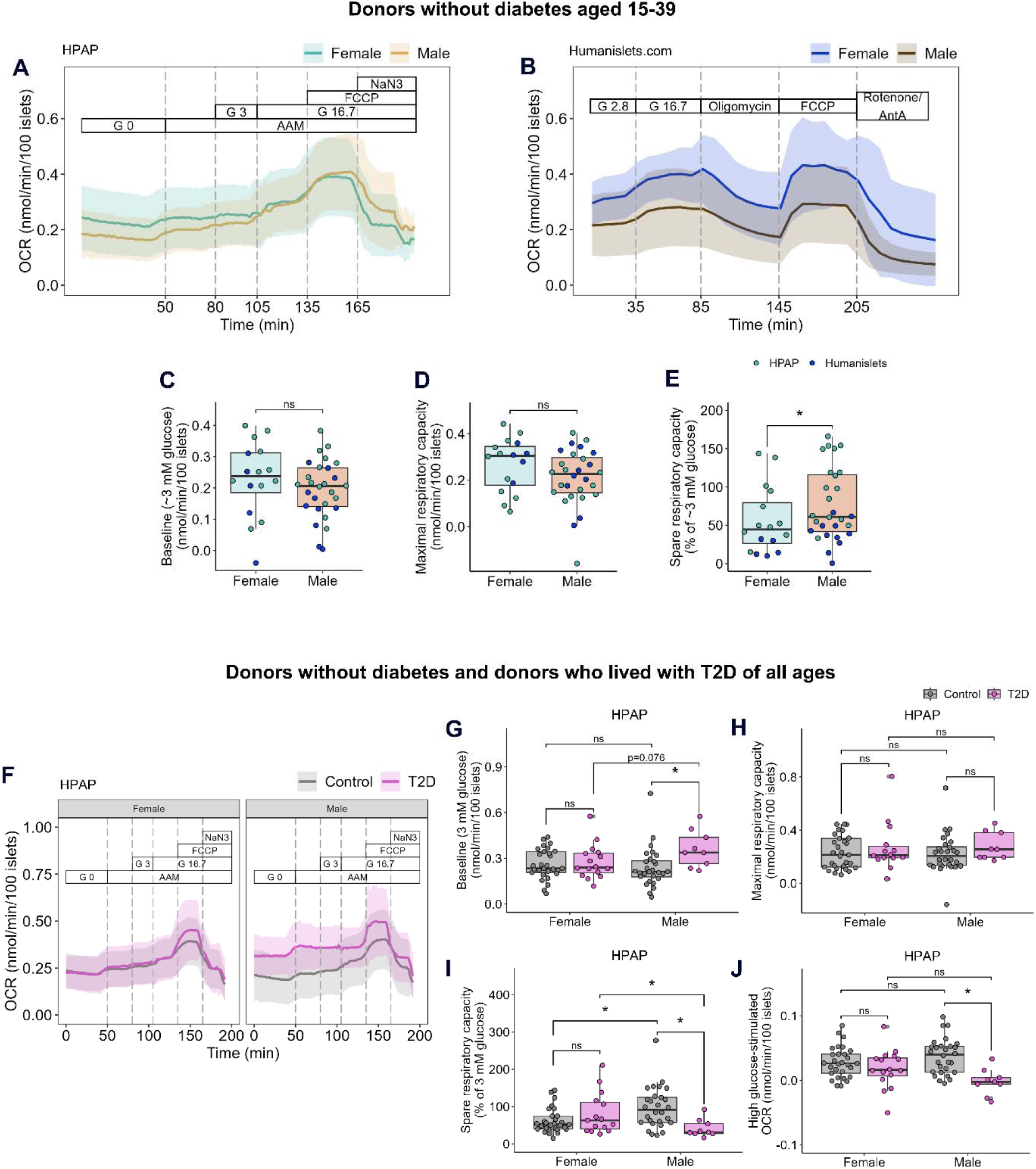
Oxygen consumption rate in whole islets from HPAP and Humanislets.com. For islets from donors without diabetes aged 15-39, average OCR ± SD tracings from HPAP (A) and Humanislets.com (B). Baseline respiration (C), maximal respiratory capacity (D), and spare respiratory capacity (E) for donors without diabetes aged 15-39 from combined HPAP and Humanislets.com data. For islets from donors of all ages without diabetes or with T2D, average OCR tracings ± SD for female and male islets from HPAP (F). Summary data for baseline OCR (G), maximal respiratory capacity (H), spare respiratory capacity (I), and high glucose-stimulated OCR (J) from donors of all ages without diabetes or with T2D, from HPAP. AAM = amino acid mix (0.44 mM alanine, 0.19 mM arginine, 0.038 mM aspartate, 0.094 mM citrulline, 0.12 mM glutamate, 0.30 mM glycine, 0.077 mM histidine, 0.094 mM isoleucine, 0.16 mM leucine, 0.37 mM lysine, 0.05 mM methionine, 0.70 mM ornithine, 0.08 mM phenylalanine, 0.35 mM proline, 0.57 mM serine, 0.27 mM threonine, 0.073 mM tryptophan, and 0.20 mM valine, 2 mM glutamine), G = glucose, FCCP = carbonyl cyanide-p-trifluoromethoxyphenylhydrazone (mitochondrial uncoupler), NaN3 = sodium azide (cytochrome oxidase inhibitor), AntA = antimycin A (complex III inhibitor). * indicates *p*<0.05, ns = not significant, *p*_≥_0.05.

We expanded our analysis to include donors of all ages, including donors who lived with T2D, focusing on the HPAP dataset due to the small number of donors who lived with T2D in the Humanislets.com dataset (Figure 6F-J). In male islets, T2D was associated with a strong impairment of mitochondrial function. Strikingly, male islets from donors with T2D showed a drastic increase in basal OCR that was not observed in female islets (Figure 6G), with no significant sex difference in maximal respiratory capacity (Figure 6H). In male islets, but not female islets, spare respiratory capacity was also significantly decreased with T2D (Figure 6I), driven by the increase in basal respiration (Figure 6J). Male islets from donors who lived with T2D also demonstrated a significantly impaired ability to increase OCR in response to high glucose; a finding we did not reproduce in female islets (Figure 6J). These T2D-associated defects in spare respiratory capacity and high glucose-stimulated respiration were also observed specifically in male islets in age-matched data, although the increase in basal respiration was no longer significant (Figure S8^41^). Thus, our data indicate that T2D is associated with fewer perturbations in islet mitochondrial parameters in females compared with males.

### Biological sex did not significantly affect intracellular calcium in whole islets

Intracellular calcium triggers insulin exocytosis ^70^. There were no significant differences in intracellular calcium following stimulation with the amino acid mixture, low glucose (3 mM), high glucose (16.7 mM), or KCl (30 mM) between the sexes, regardless of age or diabetes status (Figure S9A-E; S10A-E^41^).

### T2D is associated with impaired hormone secretion in response to stimuli in both sexes

Dynamic insulin secretion protocols were different with respect to stimulus and exposure time across all three sites. Among donors without diabetes aged 15-39, no statistically significant sex differences were observed in insulin secretion in either UPenn or Vanderbilt data (Figure S11A-L^41^). In the same age range, perifusion results from Humanislets.com showed higher insulin secretion from male islets compared to female islets (Figure S11M-Q^41^). We next considered donors of all ages and focused on the UPenn and Humanislets.com perifusions (Figure 7A-B), as these data were normalized to islet number rather than IEQ and used the same basal glucose concentration (3 mM), though we note amino acids were present in the UPenn perifusions. Combining these datasets, we found that T2D was associated with decreased insulin secretion at 3 mM glucose in males, a trend that was not significant in females (Figure 7C). In both sexes, T2D was associated with decreased insulin secretion in response to high glucose (15 or 16.7 mM, Figure 7D) and 30 mM KCl (Figure 7E). The stimulation index was decreased with T2D in both sexes (Figure 7F). These trends were unchanged when age-matched data were analyzed (Figure S12^41^). When datasets were analyzed individually, there was high variability within each dataset, reflecting the heterogeneity of nutrient-induced insulin secretion in human islets ^45,71,72^, but the T2D-related impairment was apparent in most parameters (Figure S13-14^41^). Overall, these data indicate that T2D was associated with impairment of high glucose-induced insulin secretion in both sexes.

**Figure 7.**
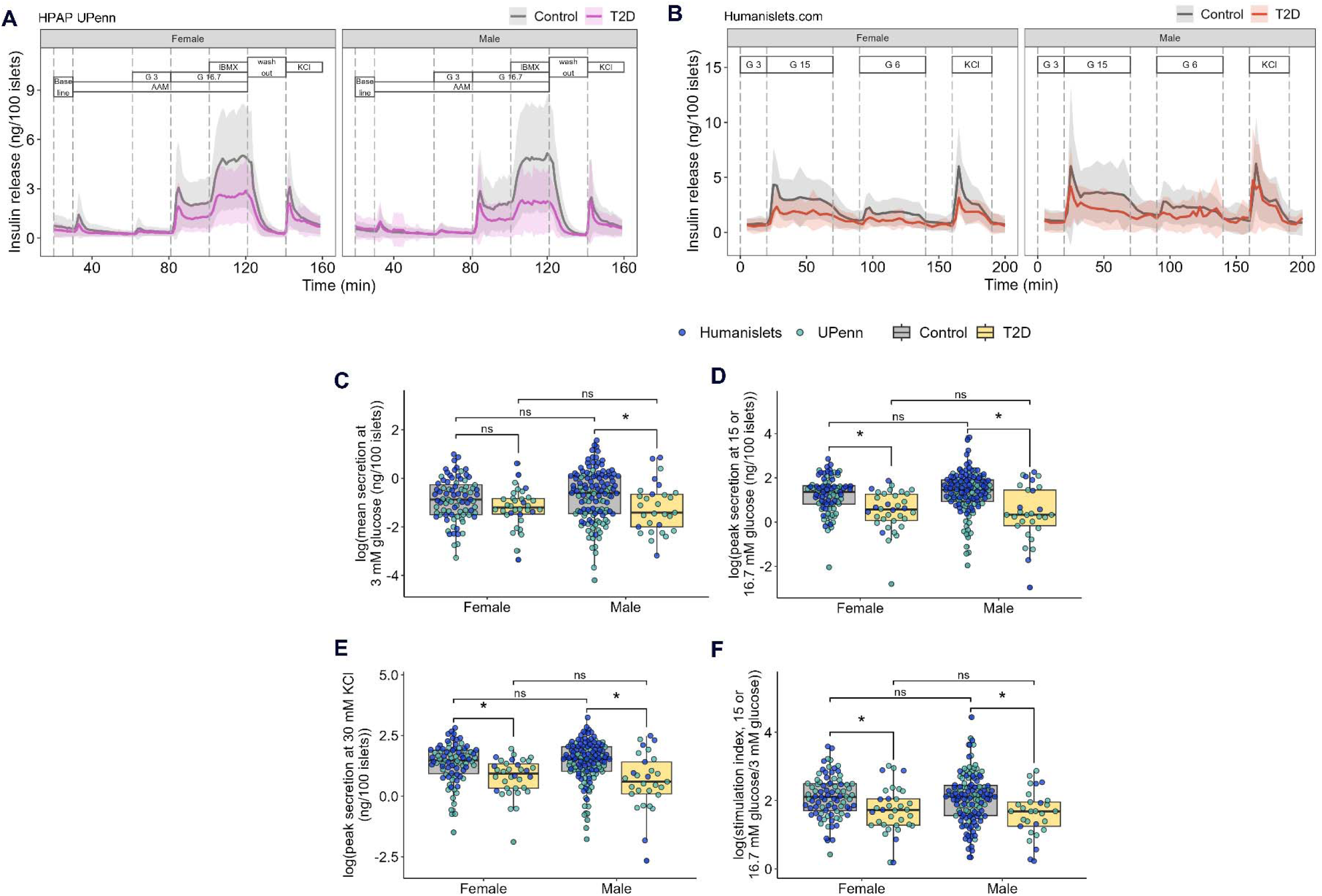
HPAP UPenn and Humanislets.com dynamic insulin secretion data following glucose stimulation for islets from donors without diabetes and donors with T2D of all ages. Perifusion curves for HPAP UPenn (A) and Humanislets.com (B) data. Average insulin secretion at 3 mM glucose for combined HPAP UPenn (AAM also present) and Humanislets.com data (C). Peak insulin secretion at high glucose for combined HPAP UPenn (16.7 mM glucose plus AAM) and Humanislets.com data (15 mM glucose) (D). Peak insulin secretion at 30 mM KCl for combined HPAP UPenn and Humanislets.com data (E). Stimulation index calculated as peak high glucose-stimulated secretion divided by average secretion at 3 mM glucose ± AAM for combined HPAP UPenn and Humanislets.com data (F). AAM = amino acid mix (0.44 mM alanine, 0.19 mM arginine, 0.038 mM aspartate, 0.094 mM citrulline, 0.12 mM glutamate, 0.30 mM glycine, 0.077 mM histidine, 0.094 mM isoleucine, 0.16 mM leucine, 0.37 mM lysine, 0.05 mM methionine, 0.70 mM ornithine, 0.08 mM phenylalanine, 0.35 mM proline, 0.57 mM serine, 0.27 mM threonine, 0.073 mM tryptophan, and 0.20 mM valine, 2 mM glutamine). G = glucose. IBMX = 3-isobutyl-1-methylxanthine at 0.1 mM. Perifusion curves show mean ± SD. * indicates *p*<0.05, ns = not significant, *p*_≥_0.05.

Humanislets.com was the only database with data on insulin secretion in response to leucine and a fatty acid mixture. Consistent with results from the glucose-focused perifusion in this donor pool (Figure S11N, S13N^41^), among donors without diabetes, male islets secreted more insulin than female islets at baseline and in response to leucine or fatty acids in both the 15-39 age bracket and across all ages (Figures S15-16^41^).

Perifusion experiments at UPenn and Vanderbilt also measured glucagon secretion and content. Among donors without diabetes aged 15-39, there were no significant sex differences in glucagon secretion or glucagon content in either dataset (Figure S17^41^). Among donors of all ages (Figure 18A-J), UPenn data showed that T2D was associated with a general decrease in glucagon secretion in female, but not male islets, in terms of basal (Figure S18C^41^), AAM-stimulated (Figure S18E^41^), and KCl-stimulated secretion (Figure S18I^41^), but this was not apparent in the Vanderbilt data (Figure S18D, F, J^41^). In both sexes, T2D was associated with decreased islet glucagon content in the UPenn data (Figure S18K^41^) but not the Vanderbilt data (Figure S18L^41^). These trends were retained when age-matched data were used (Figure S19^41^). Therefore, glucagon measurements during the perifusion studies at UPenn suggest that T2D is associated with greater abnormalities in alpha-cell glucagon secretion in female islets than male islets, but this was not observed in the Vanderbilt data.

## Discussion

The goals of the present study were to (1) identify sex differences in human islet characteristics in the non-T2D state and (2) assess how these characteristics were altered in female and male donors who lived with T2D. By performing integrated analysis of several outcomes and using two independent databases, we discovered robust sex differences across molecular, cellular, and functional phenotypes in both baseline and T2D contexts. Importantly, T2D was associated with distinct changes in gene and protein expression in female islets compared to male islets. This suggests sex-dependent mechanisms of pathogenesis, demonstrating the importance of considering biological sex in developing prevention and treatment strategies for T2D.

One key finding from our work was that female islets may be less susceptible to T2D-associated mitochondrial impairments compared to male islets. Mitochondrial respiration is not only central to GSIS ^70,73^, but also predicts transplantation success ^74–76^. Among young donors without diabetes, mitochondrial pathways were among the top female-biased pathways, consistent with prior studies in humans ^35^ and rats ^77^. While lower spare respiratory capacity was observed in female islets from young donors without diabetes, this was not associated with defects in insulin secretion. Additionally, although T2D was associated with a female-specific decrease in mitochondrial gene and protein expression, in line with a previous report ^35^, this did not translate to a functional impairment in mitochondrial respiration in islets from female donors with T2D. A potential explanation for the absence of mitochondrial impairment in female islets is the higher baseline expression of these genes, which may provide greater capacity for adaptation, possibly contributing to the lower risk of T2D ^1–4^ and later age of diagnosis ^78^ in women than in men.

Another prominent finding from our analysis was that ribosomal pathways were among the top gene and protein pathways enriched in healthy female islets. A female bias in protein synthesis pathways has been observed in several previous studies in humans ^29,33,35^, and elevated expression of ribosomal proteins in females has been found across species, including rodent islets ^28,29^ and most *Drosophila* cells ^79^. The female-biased enrichment of pathways related to both protein synthesis and ATP generation may support the energy-intensive process of synthesizing insulin ^80^. While our analysis revealed no consistent sex bias in insulin content or insulin secretion to accompany this higher expression of genes related to protein synthesis and mitochondrial function, this lack of effect may be due to the lower beta-cell proportion in female islets. Further work will also be needed to understand why T2D was associated with a positive enrichment of cytosolic ribosome genes in male islets. While these pathways may be expected to be beneficial under conditions of high insulin demand, as in T2D, it is possible that male islets have insufficient mitochondrial activity to support increased levels of protein synthesis.

We further show that alpha-cells may play a greater role in supporting female islets under physiological conditions and may be more perturbed under T2D conditions. Consistent with previous analyses in human islets ^43,71^, we confirmed that female islets had higher proportions of alpha-cells and lower proportions of beta-cells and non-alpha, non-beta endocrine cells compared to male islets. Importantly, increased alpha-cell proportion in females was observed in both the HPAP dataset and Humanislets.com datasets, which used vastly different methods to estimate this parameter. The HPAP dataset used an antibody-based CyToF method to obtain counts, which carries the caveat that islets must first be dispersed to single cells, possibly leading to losses of more fragile cell populations. The Humanislets.com dataset estimated cell type proportions based on marker protein expression in whole islet proteomics; these results may be affected by differences in cell identity. A female bias for increased alpha-cell proportion has also been observed in mice ^32^, in which alpha-cells support beta-cell function by paracrine signaling ^81–84^. Under high-fat diet stress, the greater alpha-cell proportion in females may promote beneficial intra-islet structural rearrangements that preserve beta-cell calcium dynamics ^32^. Because alpha-cell proportion further increases in female islets during pregnancy, this suggests these cells play an important role during times of additional metabolic stress ^85^. An important role for alpha-cells in supporting female metabolic health is further indicated by our finding that T2D-related changes in alpha-cell gene expression were greater in females compared to males. Indeed, the HPAP UPenn perifusion data suggested that T2D may be associated with greater changes in alpha-cell function in females than males. Together, these results highlight the importance of characterizing potential sex differences in alpha-cells, which have been understudied in the literature.

Despite multiple sex differences in islet gene and protein expression, and in cellular processes, perifusion data did not demonstrate consistent sex differences in insulin secretion. While prior analyses of earlier iterations of the HPAP islet perifusion data observed increased insulin secretion in female islets compared to male islets ^29,33,34^, inclusion of new donors shows the levels of heterogeneity were too high to detect a statistical sex difference. This is consistent with a recent analysis of insulin secretion data from 576 donors from an independent donor pool showing high heterogeneity and no measurable impact of donor sex on insulin secretion patterns^72^. However, humans demonstrate sex differences in insulin secretion *in vivo* ^86^, and acute assays in isolated human donor islets may not capture chronic changes in islet physiology before, during, and after T2D onset. More rigorous investigation of clinical data, with stratification and analysis by sex, will be required to determine how sex differences in islets at the molecular and cellular level contribute to T2D pathogenesis and treatment outcomes. Additional clinical characteristics may also affect islet-level outcomes, including current medication and potential history of gestational diabetes. Although some medication history was available in HPAP metadata, it was insufficient to perform stratified analysis. Gestational diabetes drastically increases the risk of future T2D ^87^ and is likely relevant to sex-specific disease characteristics. While there was no record of gestational diabetes among the female donors included in this study, this may be an avenue for future investigation. More work will also be needed to capture the effects of genetic ancestry on islet biology. The clinical features of T2D, including BMI at onset ^88–93^, clinical presentation ^94–97^, and genetic risk ^98–100^, can vary widely across geography and genetic ancestry. Human islet characteristics, including insulin secretion ^71,101^ and gene expression ^35^ also vary by race. Therefore, while we illustrate sex-specific differences in islet physiology, we also acknowledge that this is only a first step towards improving health equity.

In conclusion, by systematic sex-based analysis of multiple data types, we identified key sex differences in composition, gene expression, and function among donors without diabetes. These differences likely contribute to sex differences in the function of islets from donors who lived with T2D, as well as potentially sex-dependent pathways of T2D onset. Elucidating these sex-dependent disease progression pathways will be critical for developing therapeutic interventions that are effective in both sexes.

## Acknowledgements

Data from this article were downloaded from publicly available databases. HPAP data were obtained from PancDB, repository of the HPAP Database, consortia under Human Islet Research Network (RRID:SCR_014393, https://hpap.pmacs.upenn.edu/) (NIH grant numbers UC4-DK112217 and UC4-DK112232, RRID:SCR_016202). Humanislets.com data and analyses were obtained from Humanislets.com, funded by the Canadian Institutes of Health Research, JDRF Canada, and Diabetes Canada (5-SRA-2021-1149-S-B/TG 179092) with data from islets isolated by the Alberta Diabetes Institute IsletCore with the support of the Human Organ Procurement and Exchange (HOPE) program, Trillium Gift of Life Network (TGLN). We thank members of the Rideout and Johnson labs for valuable feedback and advice. We thank members of the Doliba, MacDonald and Pepper labs for clarification regarding HPAP and Humanislets.com datasets. We acknowledge that our research takes place on the traditional, ancestral, and unceded territory of the Musqueam people, and Treaty 6, 7 and 8 territories, a traditional gathering place for diverse Indigenous peoples including the Cree, Blackfoot, Métis, Nakota Sioux, Iroquois, Dene, Ojibway/ Saulteaux/Anishinaabe, Inuit, and many others.

## Data availability

Original data generated and analyzed during this study are included in this published article or in the data repositories listed in References.

