## Supplemental figures and table legends for "Biological sex affects human islet gene expression and mitochondrial function in type 2 diabetes"

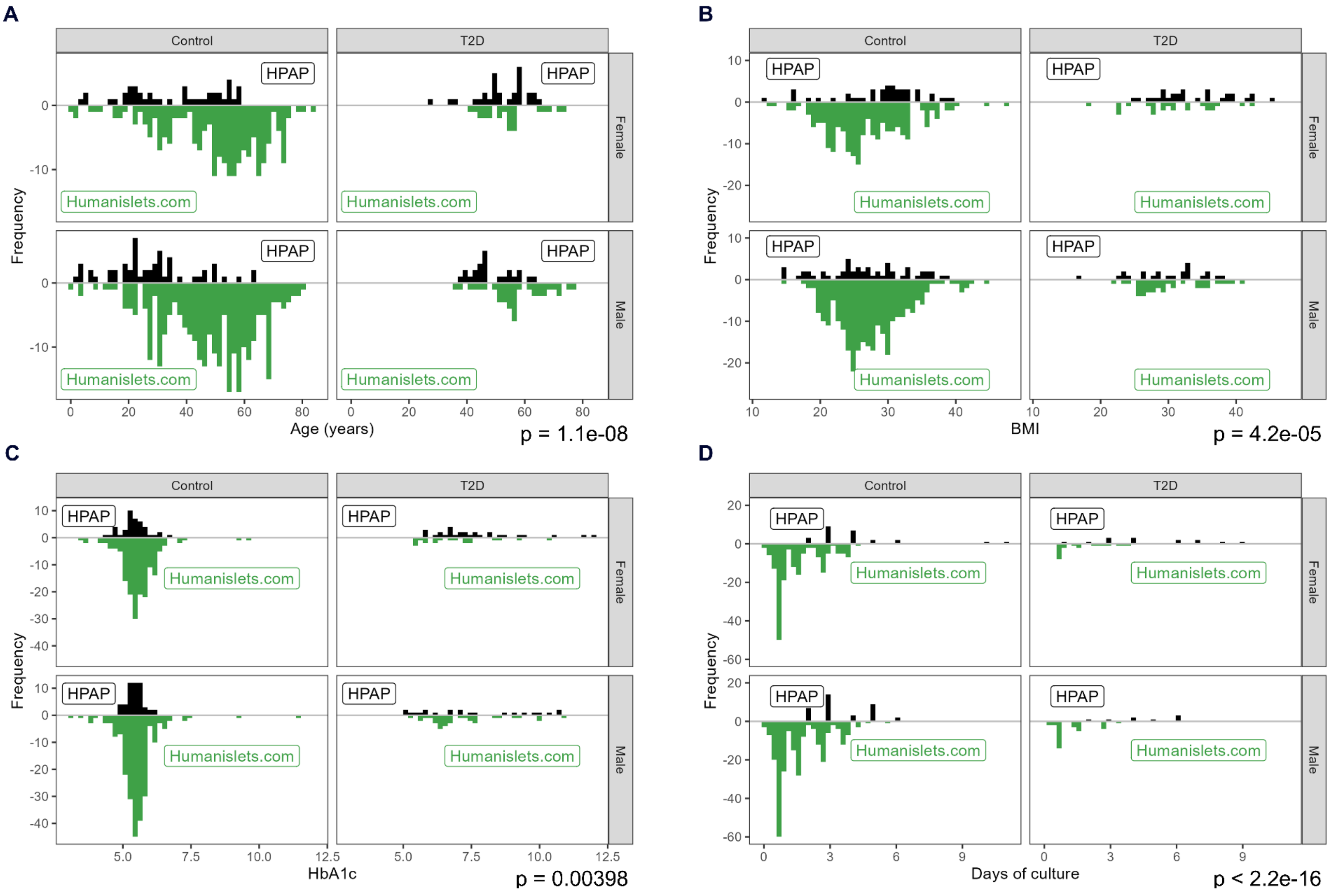

**Figure S1. Metadata distributions in HPAP and Humanislets.com datasets.**

Distribution of age (A), BMI (B), HbA1c (C), and culture duration (D). Culture duration was not available for all donors and only available data are shown. Distributions were compared by Kolgomorov-Smirnov test and *p*-values are shown in figure.

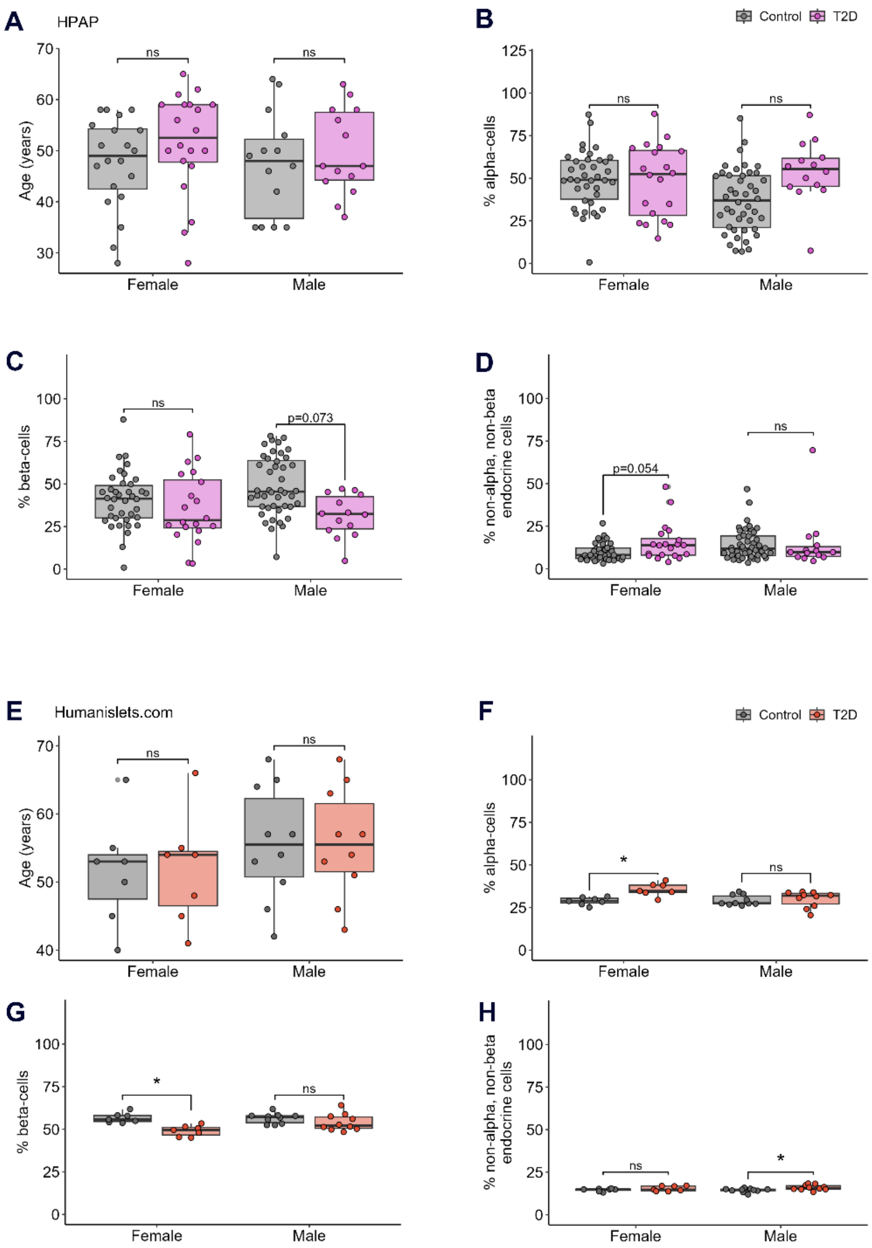


**Figure S2. Islet endocrine cell proportions in age-matched data.**

Donors who lived with T2D were age-matched with donors without T2D within each sex. Proportions were determined by CyToF in the HPAP dataset or deconvolution from whole islet proteomics in the Humanislets.com dataset. Donor ages in age-matched dataset (A), alpha-cell proportion (B), beta-cell proportion (C), and non-alpha, non-beta cell proportion (D) in age-matched HPAP data. Donor ages in age-matched dataset (E), alpha-cell proportion (F), beta-cell proportion (G), and non-alpha, non-beta cell proportion (H) in age-matched Humanislets.com data. * indicates $p<0.05$, ns = not significant, $p\geq0.05$.

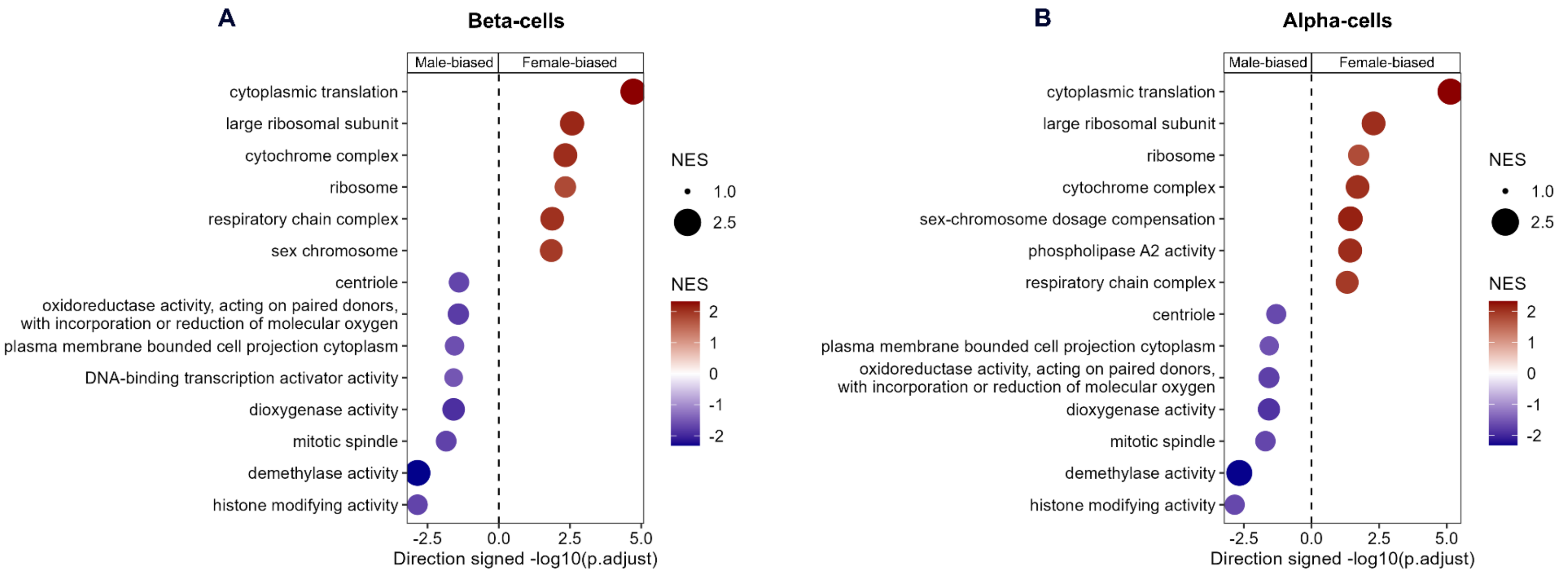


**Figure S3. Pseudobulk beta-cell and alpha-cell gene expression in islets from female versus male donors without diabetes aged 15-39.**

Significantly altered pathways according to sex in GSEA analysis in HPAP pseudobulk beta-cell scRNAseq data (A) and HPAP pseudobulk alpha-cell scRNAseq data (B). n = 7 female, 17 male. NES = normalized enrichment score.

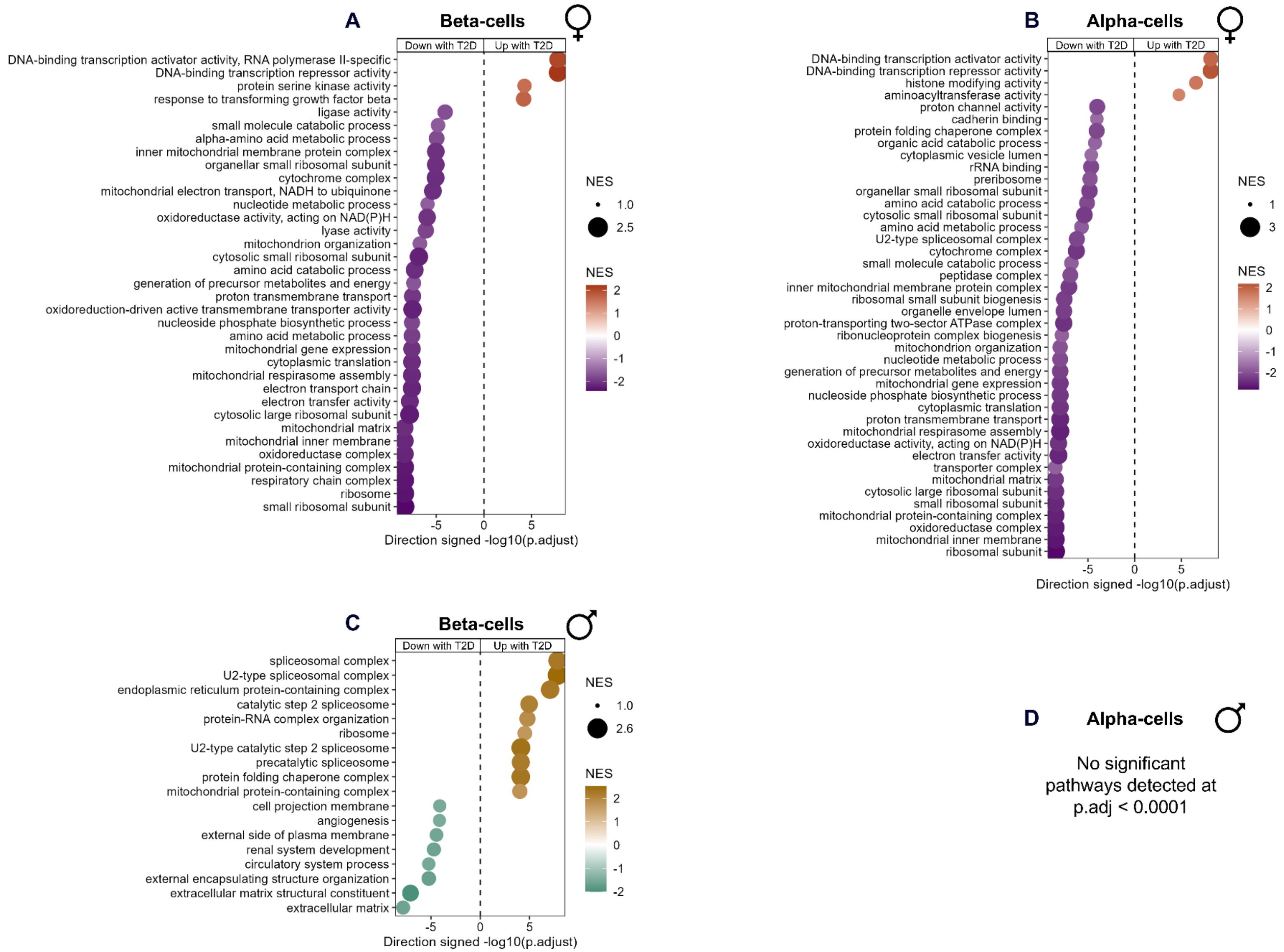
A
Beta-cells
Down with T2D
Up with T2D
DNA-binding transcription activator activity, RNA polymerase II-specific
DNA-binding transcription repressor activity
protein serine kinase activity
response to transforming growth factor beta
ligase activity
small molecule catabolic process
alpha-amino acid metabolic process
inner mitochondrial membrane protein complex
organellar small ribosomal subunit
cytochrome complex
mitochondrial electron transport, NADH to ubiquinone
nucleotide metabolic process
oxidoreductase activity, acting on NAD(P)H
lyase activity
mitochondrion organization
cytosolic small ribosomal subunit
amino acid catabolic process
generation of precursor metabolites and energy
proton transmembrane transport
oxidoreduction-driven active transmembrane transporter activity
nucleoside phosphate biosynthetic process
amino acid metabolic process
mitochondrial gene expression
cytoplasmic translation
mitochondrial respirasome assembly
electron transport chain
electron transfer activity
cytosolic large ribosomal subunit
mitochondrial matrix
mitochondrial inner membrane
oxidoreductase complex
mitochondrial protein-containing complex
respiratory chain complex
ribosome
small ribosomal subunit
-5
0
5
Direction signed -log10(p.adjust)
NES
1.0
2.5
NES
2
1
0
-1
-2
B
Alpha-cells
Down with T2D
Up with T2D
DNA-binding transcription activator activity
DNA-binding transcription repressor activity
histone modifying activity
aminoacyltransferase activity
proton channel activity
cadherin binding
protein folding chaperone complex
organic acid catabolic process
cytoplasmic vesicle lumen
rRNA binding
preribosome
organellar small ribosomal subunit
amino acid catabolic process
cytosolic small ribosomal subunit
amino acid metabolic process
U2-type spliceosomal complex
cytochrome complex
small molecule catabolic process
peptidase complex
inner mitochondrial membrane protein complex
ribosomal small subunit biogenesis
organelle envelope lumen
proton-transporting two-sector ATPase complex
ribonucleoprotein complex biogenesis
mitochondrion organization
nucleotide metabolic process
generation of precursor metabolites and energy
mitochondrial gene expression
nucleoside phosphate biosynthetic process
cytoplasmic translation
proton transmembrane transport
mitochondrial respirasome assembly
oxidoreductase activity, acting on NAD(P)H
electron transfer activity
transporter complex
mitochondrial matrix
cytosolic large ribosomal subunit
small ribosomal subunit
mitochondrial protein-containing complex
oxidoreductase complex
mitochondrial inner membrane
ribosomal subunit
-5
0
5
Direction signed -log10(p.adjust)
NES
1
3
NES
2
1
0
-1
-2
C
Beta-cells
Down with T2D
Up with T2D
spliceosomal complex
U2-type spliceosomal complex
endoplasmic reticulum protein-containing complex
catalytic step 2 spliceosome
protein-RNA complex organization
ribosome
U2-type catalytic step 2 spliceosome
precatalytic spliceosome
protein folding chaperone complex
mitochondrial protein-containing complex
cell projection membrane
angiogenesis
external side of plasma membrane
renal system development
circulatory system process
external encapsulating structure organization
extracellular matrix structural constituent
extracellular matrix
-5
0
5
Direction signed -log10(p.adjust)
NES
1.0
2.6
NES
2
1
0
-1
-2
D
Alpha-cells
No significant
pathways detected at
p.adj < 0.0001

24

**Figure S4. Pseudobulk beta-cell and alpha-cell gene expression in islets from donors without diabetes versus donors who lived with T2D of all ages.**

Significantly altered pathways according to T2D status in GSEA analysis at $p<0.0001$ in HPAP pseudobulk beta-cell scRNAseq data (A) and HPAP pseudobulk alpha-cell scRNAseq data (B) in females. Significantly altered pathways according to T2D status in GSEA analysis at $p<0.0001$ in HPAP pseudobulk beta-cell scRNAseq data in males (C). No significant pathways at $p<0.0001$ were detected in HPAP pseudobulk alpha-cell scRNAseq data in males (D). n = 16 female control, 11 female T2D, 24 male control, and 7 male T2D donors. NES = normalized enrichment score.

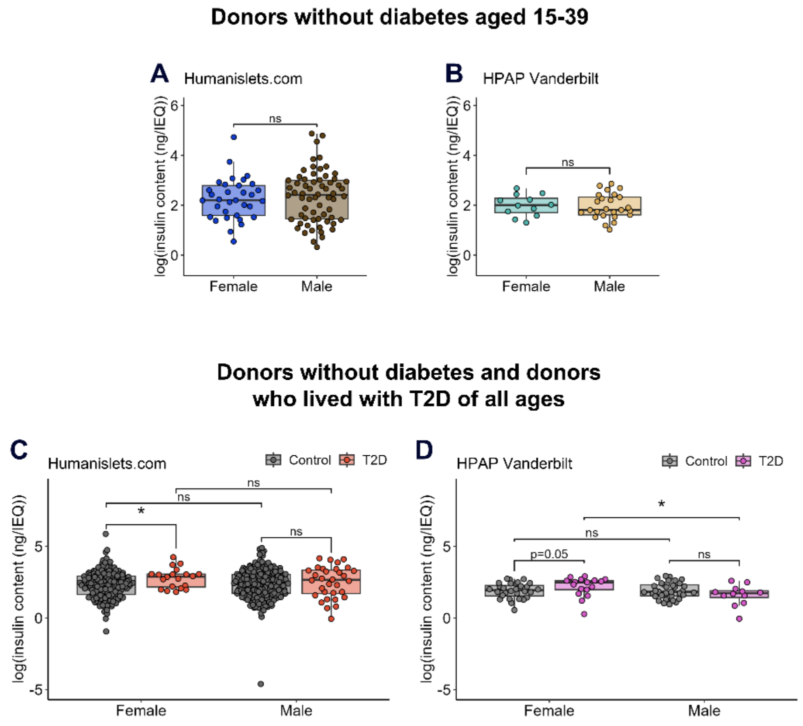


**Figure S5. Individual dataset analysis for islet insulin content.**

Insulin content from Humanislets.com (A) and HPAP Vanderbilt (B) datasets analyzed individually for donors without diabetes aged 15-39. Insulin content from Humanislets.com (C) and HPAP Vanderbilt (D) datasets analyzed individually for donors without diabetes and donors with T2D of all ages. * indicates $p<0.05$, ns = not significant, $p \geq 0.05$.

Supplement

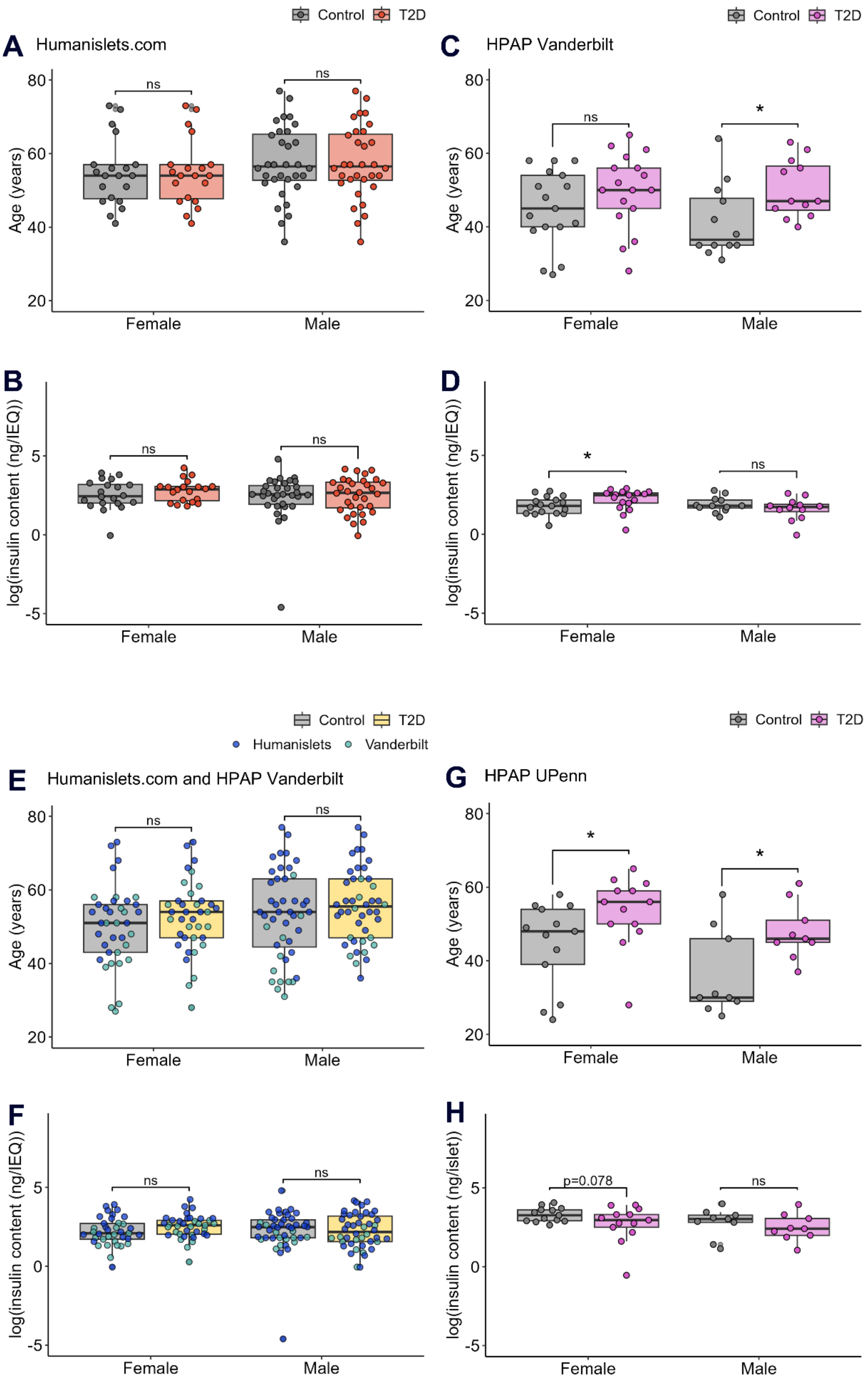


40

**Figure S6. Islet insulin content in age-matched data.**

Donors who lived with T2D were age-matched as closely as possible with donors without T2D within each sex. Donor ages in age-matched data (A) and insulin content (B) in age-matched Humanislets.com data. Donor ages in age-matched data (C) and insulin content (D) in age-matched HPAP Vanderbilt data. Donor ages in age-matched data (E) and insulin content (F) in age-matched combined Humanislets.com and HPAP Vanderbilt data. Donor ages in age-matched data (G) and insulin content (H) in age-matched HPAP UPenn data. * indicates $p<0.05$, ns = not significant, $p\geq0.05$.

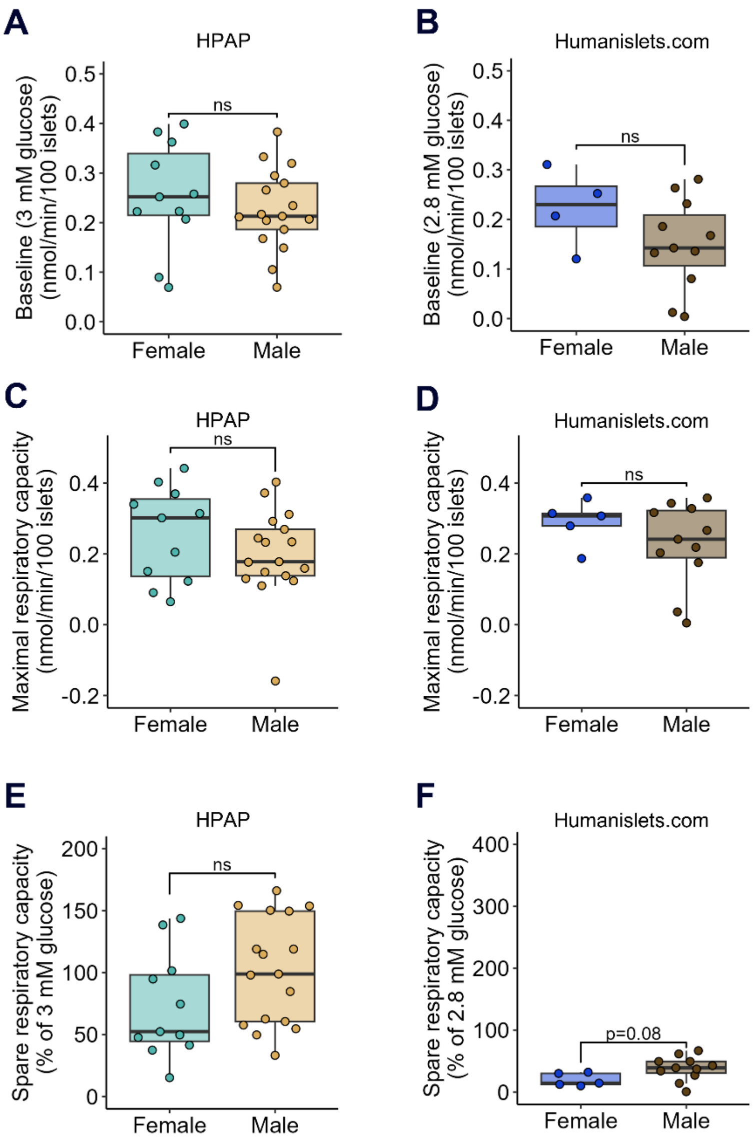


**Figure S7. Individual dataset analysis for oxygen consumption rate.**

For islets from donors without diabetes aged 15-39, baseline respiration from HPAP (A) and Humanislets.com (B), maximal respiratory capacity from HPAP (C) and Humanislets.com (D), and spare respiratory capacity from HPAP (E) and Humanislets.com (F), with datasets analyzed individually. ns = not significant, $p \geq 0.05$.

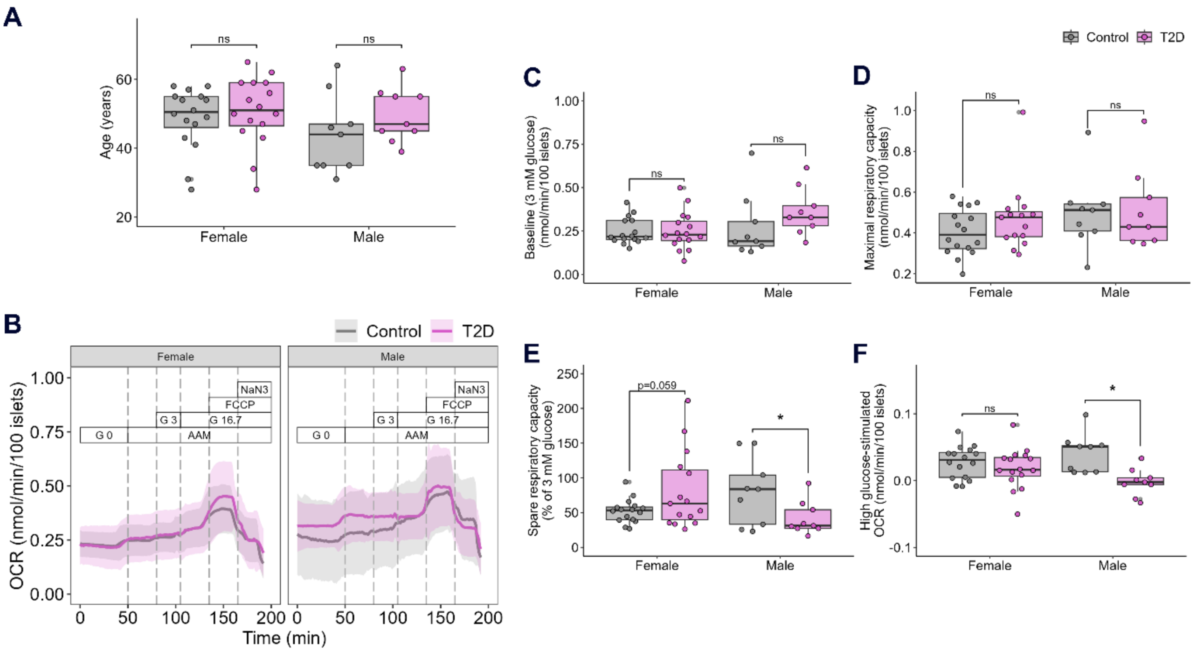


**Figure S8. HPAP oxygen consumption rate in age-matched data.**

Donors who lived with T2D were age-matched with donors without T2D within each sex. Donor ages in age-matched data (A) and average OCR tracings ± SD for female and male islets for age-matched data (B). Summary data for baseline OCR (C), maximal respiratory capacity (D), spare respiratory capacity (E), and high glucose-stimulated OCR (F) for age-matched data. AAM = amino acid mix (0.44 mM alanine, 0.19 mM arginine, 0.038 mM aspartate, 0.094 mM citrulline, 0.12 mM glutamate, 0.30 mM glycine, 0.077 mM histidine, 0.094 mM isoleucine, 0.16 mM leucine, 0.37 mM lysine, 0.05 mM methionine, 0.70 mM ornithine, 0.08 mM phenylalanine, 0.35 mM proline, 0.57 mM serine, 0.27 mM threonine, 0.073 mM tryptophan, and 0.20 mM valine, 2 mM glutamine), G 3 = 3 mM glucose, G 16.7 = 16.7 mM glucose, FCCP = carbonyl cyanide-p-trifluoromethoxyphenylhydrazone (mitochondrial uncoupler), NaN3 = sodium azide (cytochrome oxidase inhibitor), AntA = antimycin A (complex III inhibitor). * indicates $p<0.05$, ns = not significant, $p \geq 0.05$.

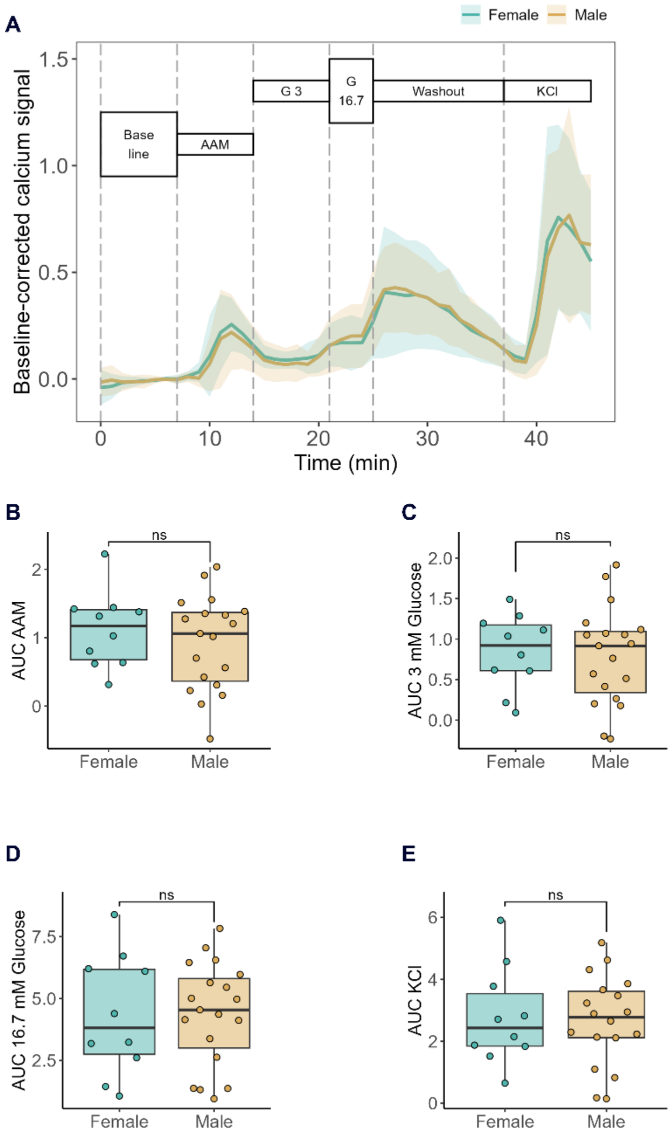


**Figure S9. HPAP intracellular calcium imaging data for islets from donors without diabetes aged 15-39.**

Average baseline-corrected intracellular calcium signal ± SD for female and male islets from donors without diabetes aged 15-39 (A). Area under the curve (AUC) data for AAM-stimulated (B), low glucose-stimulated (C), high glucose-stimulated (D), and KCl-stimulated (E) calcium signal. AAM = amino acid mix (0.44 mM alanine, 0.19 mM arginine, 0.038 mM aspartate, 0.094 mM citrulline, 0.12 mM glutamate, 0.30 mM glycine, 0.077 mM histidine, 0.094 mM isoleucine, 0.16 mM leucine, 0.37 mM lysine, 0.05 mM methionine, 0.70 mM ornithine, 0.08 mM phenylalanine, 0.35 mM proline, 0.57 mM serine, 0.27 mM threonine, 0.073 mM tryptophan, and 0.20 mM valine, 2 mM glutamine), G 3 = 3 mM glucose, G 16.7 = 16.7 mM glucose. ns = not significant, $p \geq 0.05$.

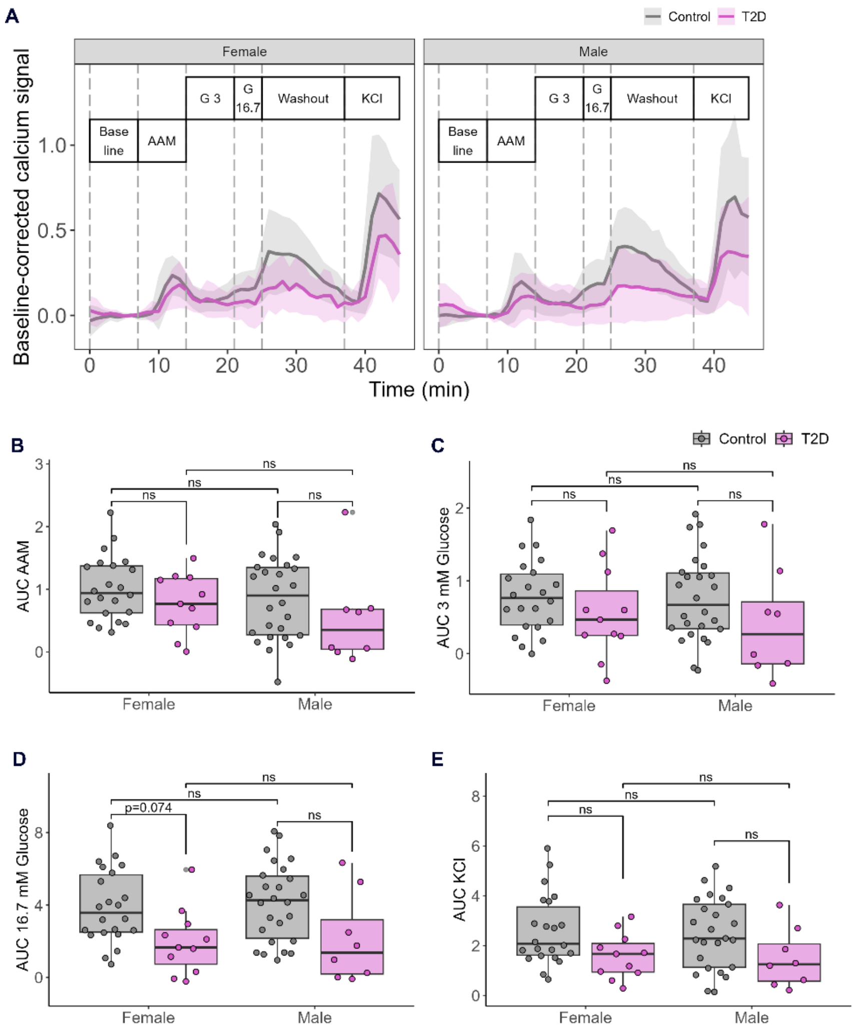


**Figure S10. HPAP intracellular calcium imaging data for islets from donors without diabetes and donors with T2D of all ages.**

Average baseline-corrected intracellular calcium signal ± SD for control and T2D, female and male islets (A). Area under the curve (AUC) data for AAM-stimulated (B), low glucose-stimulated (C), high glucose-stimulated (D), and KCl-stimulated (E) calcium signal. AAM = amino acid mix (0.44 mM alanine, 0.19 mM arginine, 0.038 mM aspartate, 0.094 mM citrulline, 0.12 mM glutamate, 0.30 mM glycine, 0.077 mM histidine, 0.094 mM isoleucine, 0.16 mM leucine, 0.37 mM lysine, 0.05 mM methionine, 0.70 mM ornithine, 0.08 mM phenylalanine, 0.35 mM proline,

0.57 mM serine, 0.27 mM threonine, 0.073 mM tryptophan, and 0.20 mM valine, 2 mM glutamine), G 3 = 3 mM glucose, G 16.7 = 16.7 mM glucose. ns = not significant, $p \geq 0.05$.

Supplement

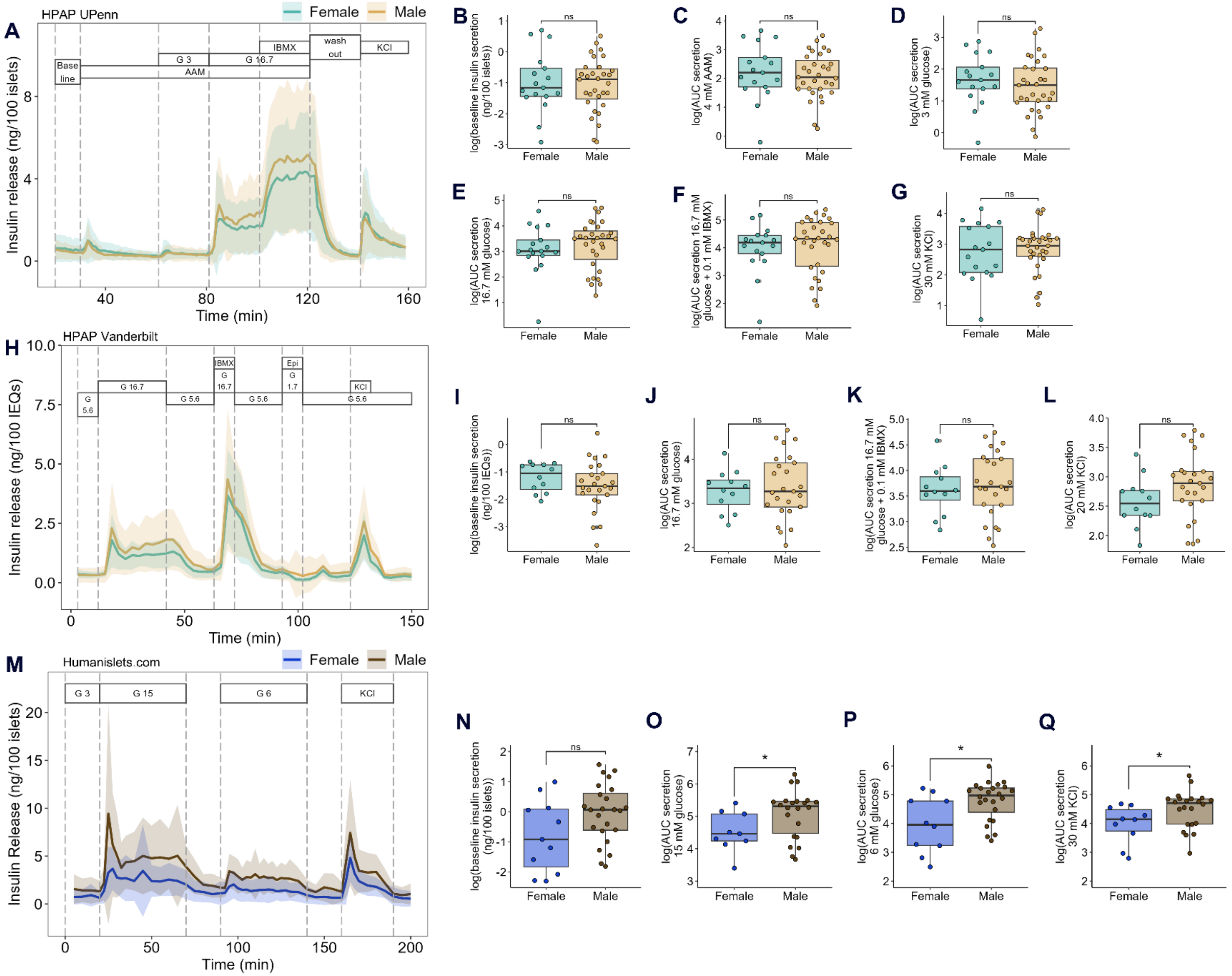
A
HPAP UPenn
Female
Male
Insulin release (ng/100 islets)
Time (min)
Base line
G 3
G 16.7
IBMX
wash out
KCl
AAM
B
C
D
E
F
G
log(baseline insulin secretion (ng/100 islets))
log(AUC secretion 4 mM AAM)
log(AUC secretion 3 mM glucose)
log(AUC secretion 16.7 mM glucose)
log(AUC secretion 16.7 mM glucose + 0.1 mM IBMX)
log(AUC secretion 30 mM KCl)
ns
H
HPAP Vanderbilt
Insulin release (ng/100 IEQs)
G 5.6
G 16.7
Epi
G 1.7
KCl
I
J
K
L
log(baseline insulin secretion (ng/100 IEQs))
log(AUC secretion 20 mM KCl)
M
Humanislets.com
Insulin Release (ng/100 islets)
G 15
G 6
N
O
P
Q
log(AUC secretion 15 mM glucose)
log(AUC secretion 6 mM glucose)
*

**Figure S11. HPAP and Humanislets.com dynamic insulin secretion data following glucose stimulation for islets from donors without diabetes aged 15-39.**

Perifusion curves (A), area under the curve for baseline (B), amino acid-induced (C), low glucose-induced (D), high glucose-induced (E), IBMX-induced (F), and KCl-induced (G) insulin secretion for HPAP UPenn data. Perifusion curves (H), baseline insulin secretion (I), area under the curve for high glucose-induced (J), IBMX-induced (K) and KCl-induced (L) insulin secretion for HPAP Vanderbilt data. Perifusion curves (M), baseline insulin secretion (N), area under the curve for high glucose-induced (O), low glucose induced (P), and KCl-induced (Q) insulin secretion for Humanislets.com data. AAM = amino acid mix (0.44 mM alanine, 0.19 mM arginine, 0.038 mM aspartate, 0.094 mM citrulline, 0.12 mM glutamate, 0.30 mM glycine, 0.077 mM histidine, 0.094 mM isoleucine, 0.16 mM leucine, 0.37 mM lysine, 0.05 mM methionine, 0.70 mM ornithine, 0.08 mM phenylalanine, 0.35 mM proline, 0.57 mM serine, 0.27 mM threonine, 0.073 mM tryptophan, and 0.20 mM valine, 2 mM glutamine), G = glucose, IBMX = 3-isobutyl-1-methylxanthine at 0.1 mM. KCl is at 30 mM for UPenn and Humanislets.com, 20 mM for Vanderbilt. Perifusion curves show mean ± SD. * indicates $p<0.05$, ns = not significant, $p \geq 0.05$.

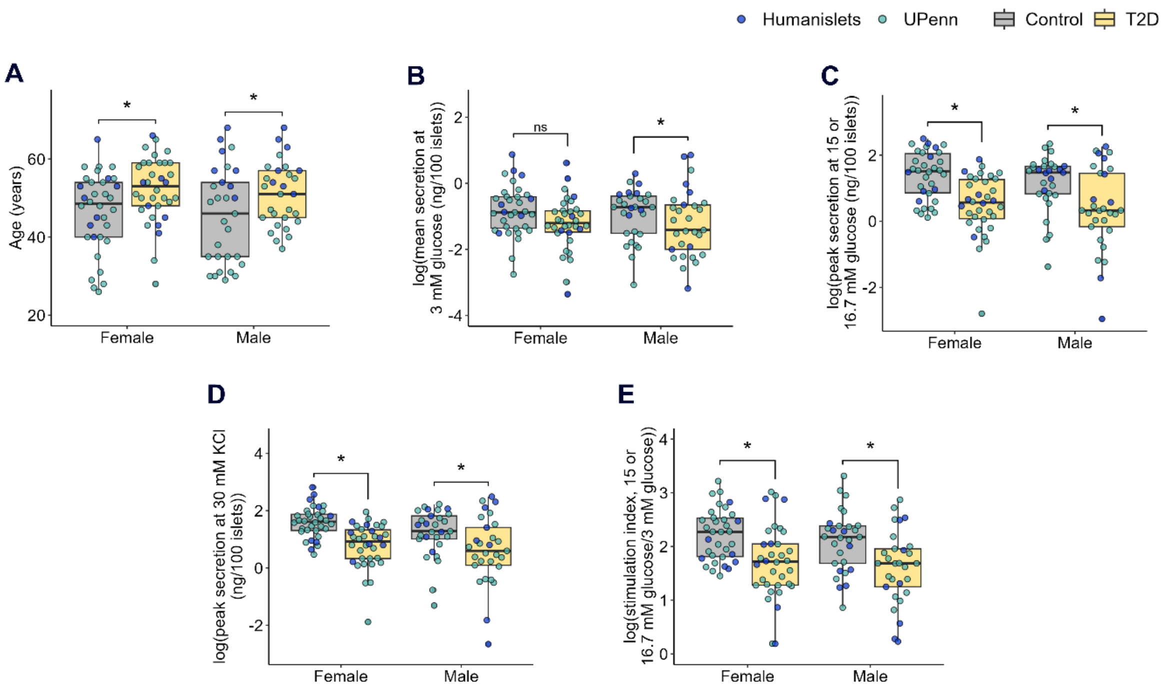


**Figure S12. Combined HPAP UPenn and Humanislets.com dynamic insulin secretion data in age-matched data.**

Donors who lived with T2D were age-matched with donors without T2D within each sex and dataset. Donor ages in age-matched data (A). Average insulin secretion at 3 mM glucose for age-matched combined HPAP UPenn (AAM also present) and Humanislets.com data (C). Peak insulin secretion at high glucose for age-matched combined HPAP UPenn (16.7 mM glucose plus AAM) and Humanislets.com data (15 mM glucose) (D). Peak insulin secretion at 30 mM KCl for age-matched combined HPAP UPenn and Humanislets.com data (E). Stimulation index calculated as peak high glucose-stimulated secretion divided by average secretion at 3 mM glucose ± AAM for age-matched combined HPAP UPenn and Humanislets.com data (F). * indicates *p*<0.05, ns = not significant, *p*≥0.05.

Supplement

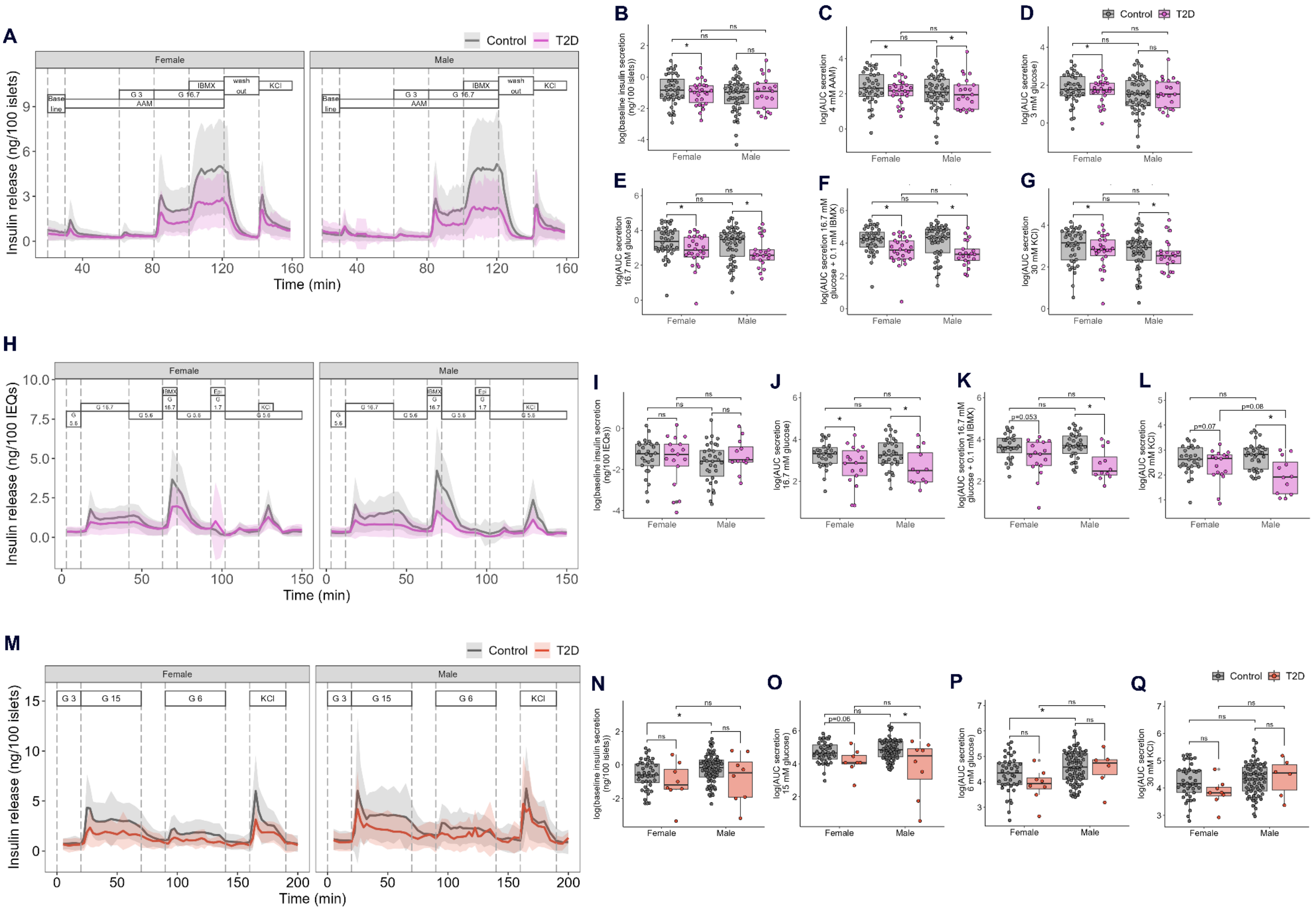
A
Control
T2D
Female
Male
Base line
G 3
AAM
G 16.7
IBMX
wash out
KCl
Insulin release (ng/100 islets)
Time (min)
B
C
D
E
F
G
log(baseline insulin secretion (ng/100 islets))
log(AUC secretion 4 mM AAM)
log(AUC secretion 3 mM glucose)
log(AUC secretion 16.7 mM glucose)
log(AUC secretion 16.7 mM glucose + 0.1 mM IBMX)
log(AUC secretion 30 mM KCl)
H
G 5.6
Epi
Insulin release (ng/100 IEQs)
I
J
K
L
log(baseline insulin secretion (ng/100 IEQs))
log(AUC secretion 20 mM KCl)
p=0.053
p=0.07
p=0.08
M
G 15
G 6
N
O
P
Q
log(AUC secretion 15 mM glucose)
log(AUC secretion 6 mM glucose)
p=0.06
ns
*

**Figure S13. Individual dataset analysis for HPAP UPenn, HPAP Vanderbilt, and Humanislets.com dynamic insulin secretion data for islets from donors without diabetes and donors with T2D of all ages.**

Perifusion curves (A), average baseline (0 mM glucose, B), and area under the curve for amino acid-induced (C), 3 mM glucose-induced (D), 16.7 mM glucose-induced (E), 16.7 mM glucose and IBMX-induced (F), and 30 mM KCl-induced (G) insulin secretion for HPAP UPenn data. Perifusion curves (H), average baseline (5.6 mM glucose, I), and area under the curve for 16.7 mM glucose-induced (J), 16.7 mM glucose and IBMX-induced (K) and 20 mM KCl-induced (L) insulin secretion for HPAP Vanderbilt data. Perifusion curves (M), average baseline (3 mM glucose, N), and area under the curve for 15 mM glucose-induced (O), 6 mM glucose-induced (P) and 30 mM KCl-induced (Q) insulin secretion for Humanislets.com data. AAM = amino acid mix (0.44 mM alanine, 0.19 mM arginine, 0.038 mM aspartate, 0.094 mM citrulline, 0.12 mM glutamate, 0.30 mM glycine, 0.077 mM histidine, 0.094 mM isoleucine, 0.16 mM leucine, 0.37 mM lysine, 0.05 mM methionine, 0.70 mM ornithine, 0.08 mM phenylalanine, 0.35 mM proline, 0.57 mM serine, 0.27 mM threonine, 0.073 mM tryptophan, and 0.20 mM valine, 2 mM glutamine), G = glucose, IBMX = 3-isobutyl-1-methylxanthine at 0.1 mM. Perifusion curves show mean ± SD. * indicates $p<0.05$, ns = not significant, $p\geq0.05$.

Supplement

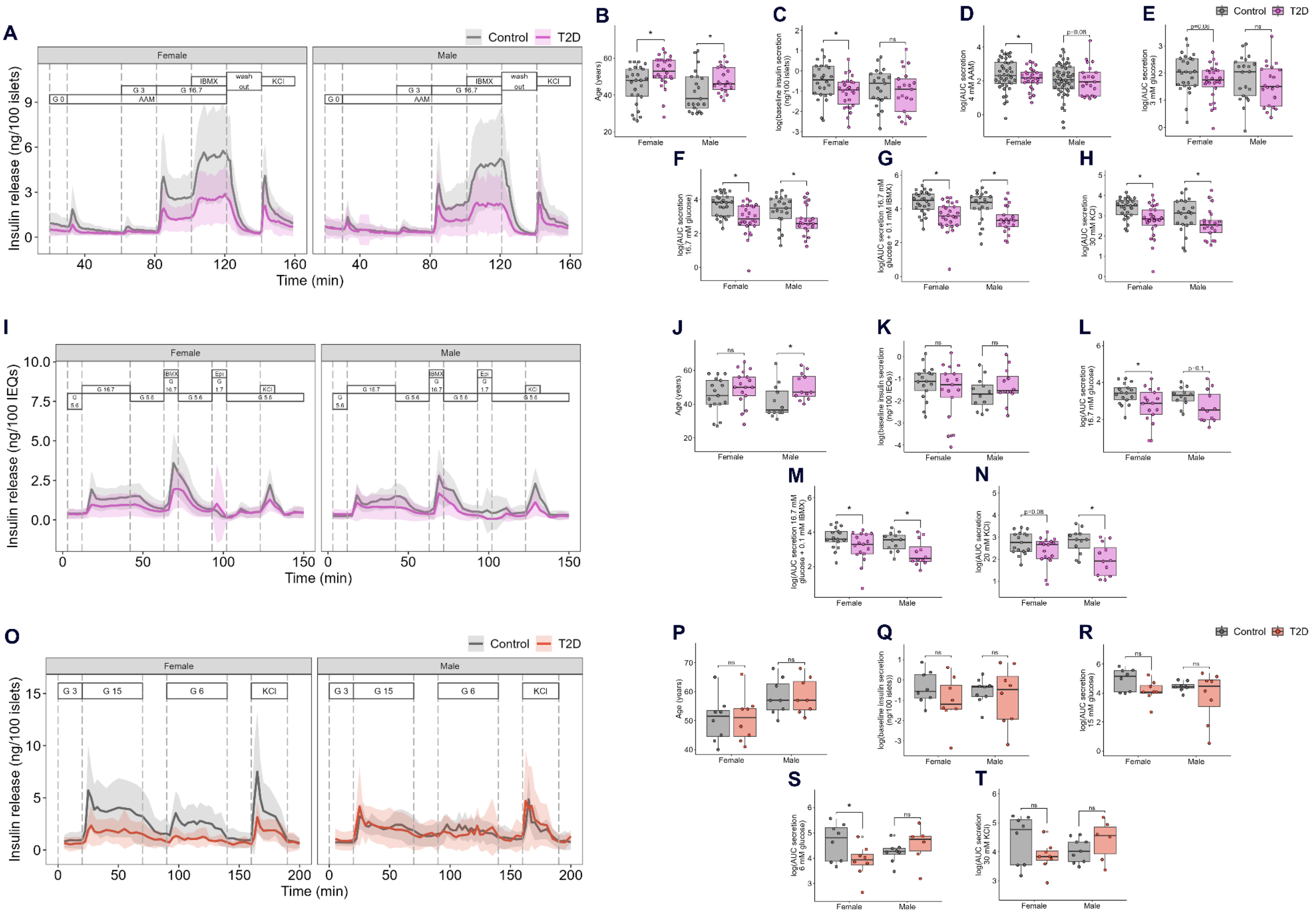
A
Control
T2D
Female
Male
G 0
G 3
AAM
G 16.7
IBMX
wash out
KCl
Insulin release (ng/100 islets)
Time (min)
B
C
D
E
F
G
H
Age (years)
log(baseline insulin secretion (ng/100 islets))
log(AUC secretion 4 mM AAM)
log(AUC secretion 3 mM glucose)
log(AUC secretion 16.7 mM glucose)
log(AUC secretion 16.7 mM glucose + 0.1 mM IBMX)
log(AUC secretion 30 mM KCl)
ns
p=0.08
I
G 5.6
Epi
Insulin release (ng/100 IEQs)
J
K
L
M
N
log(baseline insulin secretion (ng/100 IEQs))
log(AUC secretion 20 mM KCl)
p=0.1
O
G 15
G 6
P
Q
R
S
T
log(AUC secretion 15 mM glucose)
log(AUC secretion 6 mM glucose)

**Figure S14. Individual dataset analysis for HPAP UPenn, HPAP Vanderbilt, and Humanislets.com dynamic insulin secretion in age-matched data.**

Donors who lived with T2D were age-matched as closely as possible with donors without T2D within each sex. Perifusion curves (A), donor ages (B), average baseline (0 mM glucose, C), and area under the curve for amino acid-induced (D), 3 mM glucose-induced (E), 16.7 mM glucose-induced (F), 16.7 mM glucose and IBMX-induced (G), and 30 mM KCl-induced (H) insulin secretion for age-matched HPAP UPenn data. Perifusion curves (I), donor ages (J), average baseline (5.6 mM glucose, K), and area under the curve for 16.7 mM glucose-induced (L), 16.7 mM glucose and IBMX-induced (M) and 20 mM KCl-induced (N) insulin secretion for age-matched HPAP Vanderbilt data. Perifusion curves (O), donor ages (P), average baseline (3 mM glucose, Q), and area under the curve for 15 mM glucose-induced (R), 6 mM glucose-induced (S) and 30 mM KCl-induced (T) insulin secretion for age-matched Humanislets.com data. AAM = amino acid mix (0.44 mM alanine, 0.19 mM arginine, 0.038 mM aspartate, 0.094 mM citrulline, 0.12 mM glutamate, 0.30 mM glycine, 0.077 mM histidine, 0.094 mM isoleucine, 0.16 mM leucine, 0.37 mM lysine, 0.05 mM methionine, 0.70 mM ornithine, 0.08 mM phenylalanine, 0.35 mM proline, 0.57 mM serine, 0.27 mM threonine, 0.073 mM tryptophan, and 0.20 mM valine, 2 mM glutamine), G = glucose, IBMX = 3-isobutyl-1-methylxanthine at 0.1 mM. Perifusion curves show mean ± SD. * indicates $p$<0.05, ns = not significant, $p$≥0.05.

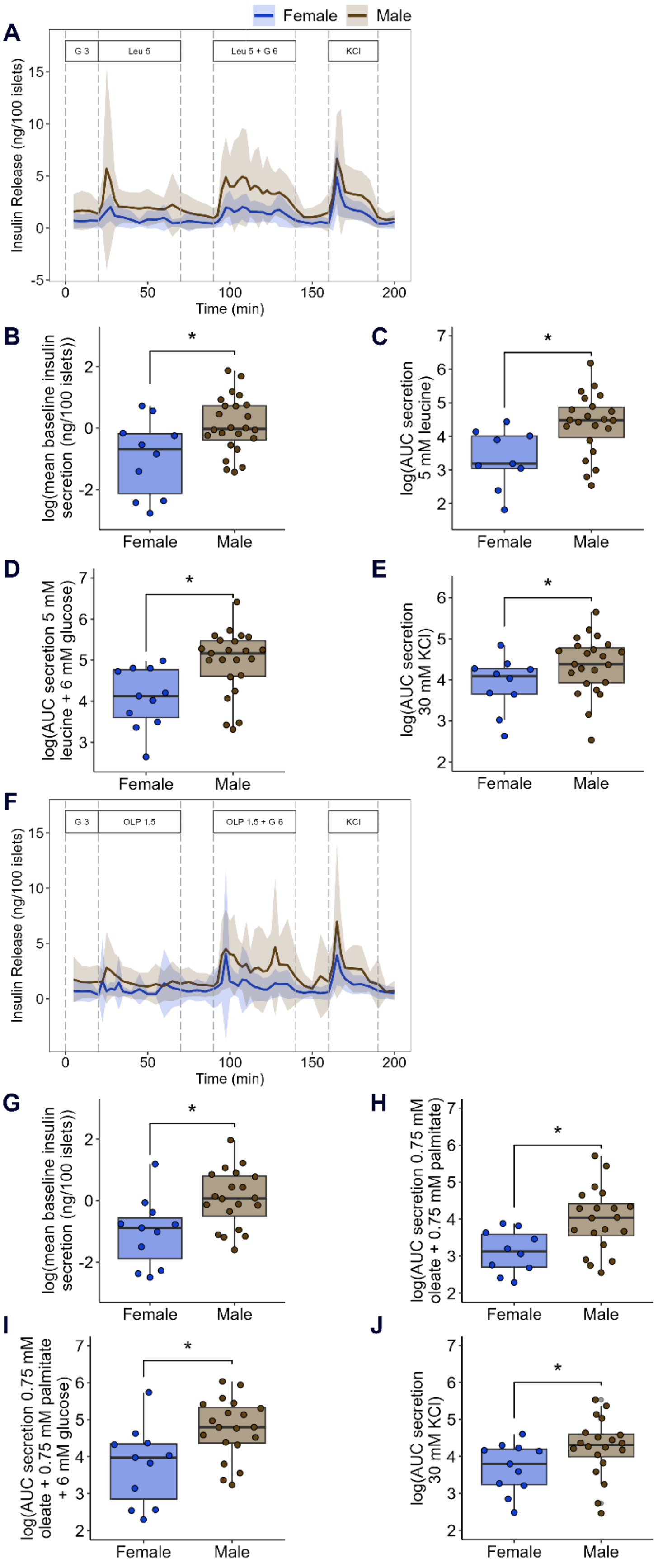
Female
Male
A
G 3
Leu 5
Leu 5 + G 6
KCl
Insulin Release (ng/100 islets)
Time (min)
B
log(mean baseline insulin secretion (ng/100 islets))
C
log(AUC secretion 5 mM leucine)
D
log(AUC secretion 5 mM leucine + 6 mM glucose)
E
log(AUC secretion 30 mM KCl)
F
G 3
OLP 1.5
OLP 1.5 + G 6
KCl
Insulin Release (ng/100 islets)
Time (min)
G
log(mean baseline insulin secretion (ng/100 islets))
H
log(AUC secretion 0.75 mM oleate + 0.75 mM palmitate)
I
log(AUC secretion 0.75 mM oleate + 0.75 mM palmitate + 6 mM glucose)
J
log(AUC secretion 30 mM KCl)
*

**Figure S15. Humanislets.com dynamic insulin secretion data following leucine or fatty acid stimulation for islets from donors without diabetes aged 15-39.**

Perifusion curves (A) and summary statistics for baseline insulin secretion (B), leucine-induced insulin secretion (C), nutrient combination-induced insulin secretion (D), and KCl-induced insulin secretion (E) for leucine perifusion. Perifusion curves (F) and summary statistics for baseline insulin secretion (G), oleate/palmitate-induced insulin secretion (H), nutrient combination-induced insulin secretion (I), and KCl-induced insulin secretion (J) for oleate + palmitate perifusion. Perifusion curves show mean ± SD. * indicates $p<0.05$, ns = not significant, $p\geq0.05$.

Supplement

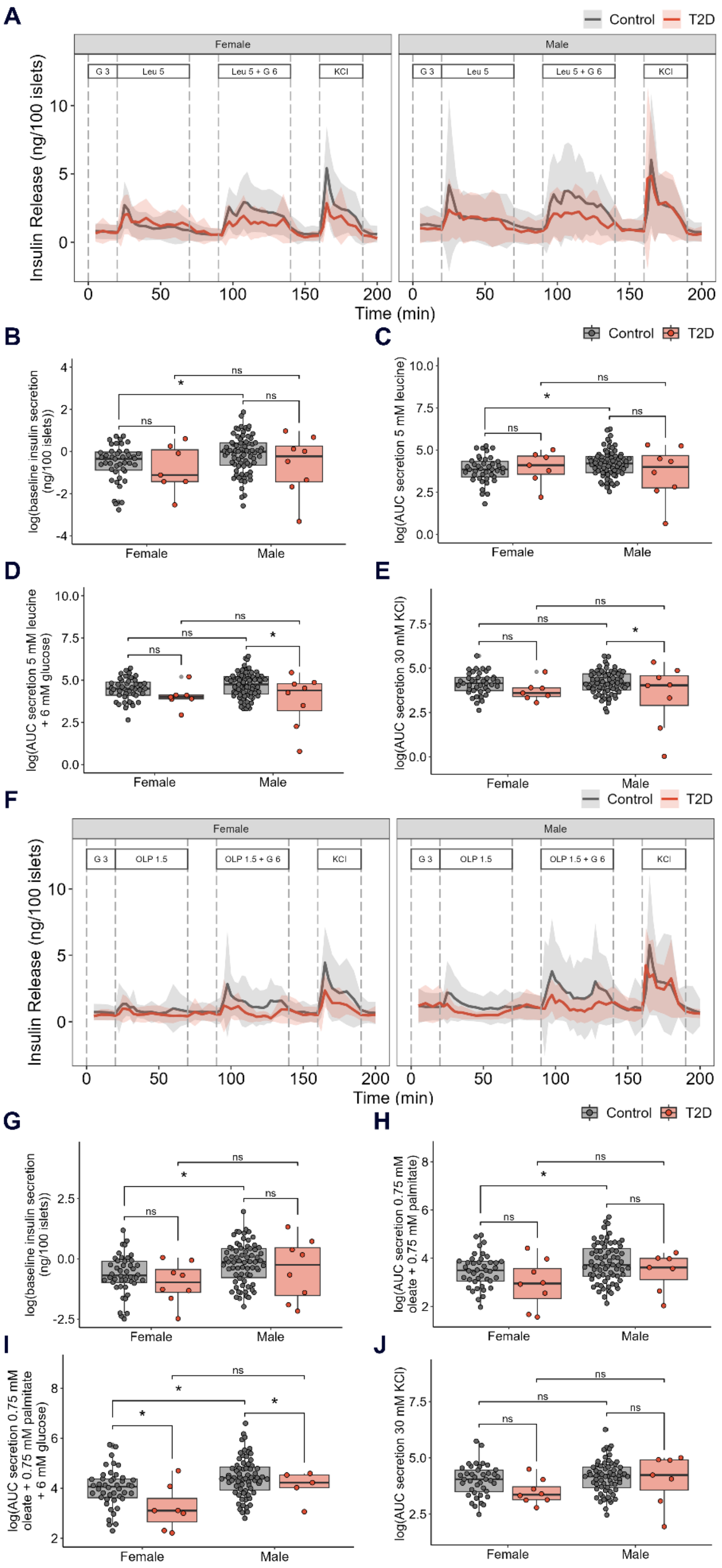
A
Control
T2D
Female
Male
G 3
Leu 5
Leu 5 + G 6
KCl
Insulin Release (ng/100 islets)
Time (min)
B
C
log(baseline insulin secretion (ng/100 islets))
log(AUC secretion 5 mM leucine)
D
E
log(AUC secretion 5 mM leucine + 6 mM glucose)
log(AUC secretion 30 mM KCl)
F
OLP 1.5
OLP 1.5 + G 6
G
H
log(AUC secretion 0.75 mM oleate + 0.75 mM palmitate)
I
J
log(AUC secretion 0.75 mM palmitate oleate + 0.75 mM palmitate + 6 mM glucose)
ns
*

**Figure S16. Humanislets.com dynamic insulin secretion data following leucine or fatty acid stimulation for islets from donors without diabetes and donors with T2D of all ages.** Perifusion curves (A) and summary statistics for baseline insulin secretion (B), leucine-induced insulin secretion (C), nutrient combination-induced insulin secretion (D), and KCl-induced insulin secretion (E) for leucine perifusion. Perifusion curves (F) and summary statistics for baseline insulin secretion (G), oleate/palmitate-induced insulin secretion (H), nutrient combination-induced insulin secretion (I), and KCl-induced insulin secretion (J) for oleate + palmitate perifusion. Perifusion curves show mean ± SD. * indicates $p<0.05$, ns = not significant, $p \geq 0.05$.

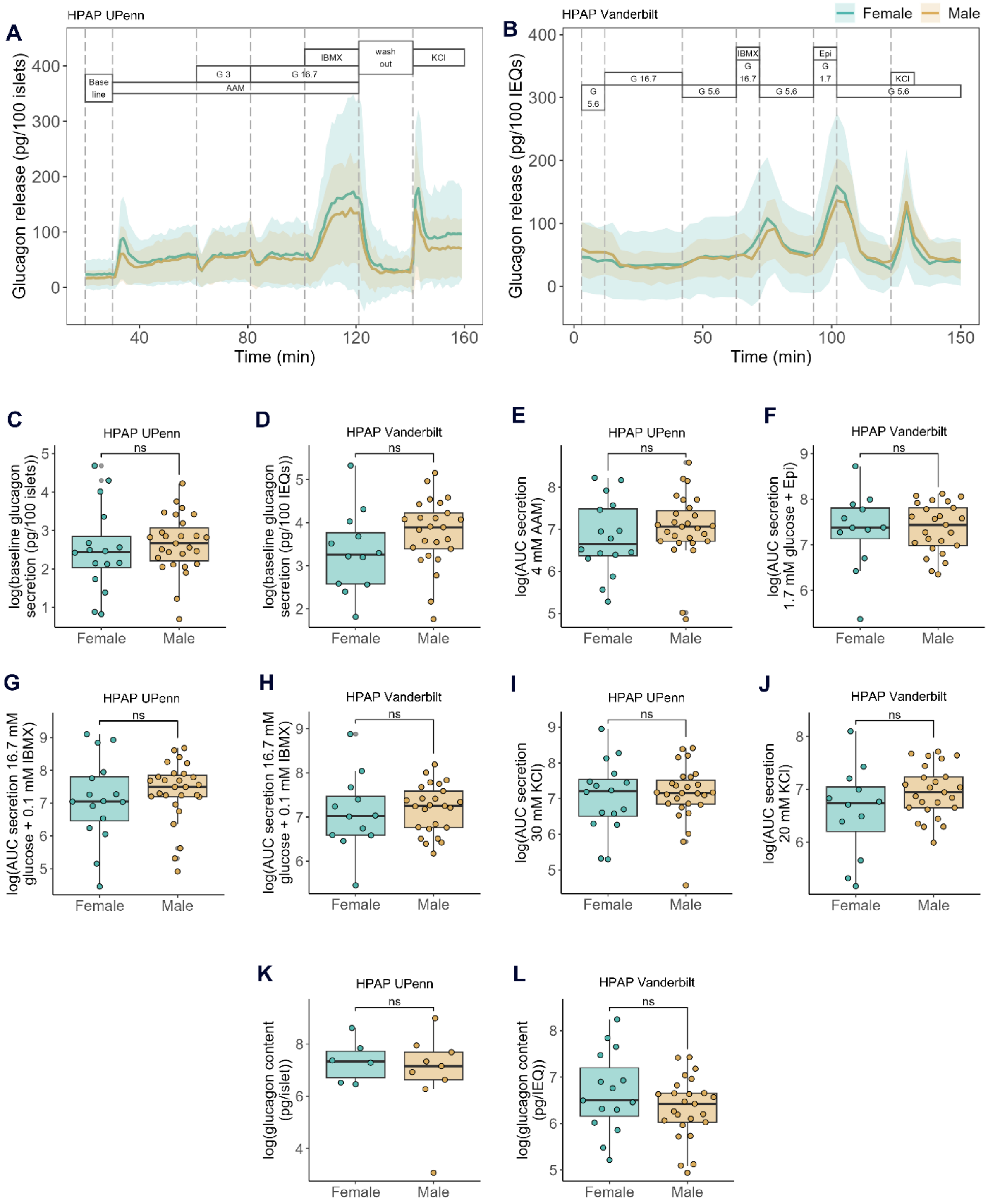


**Figure S17. HPAP dynamic glucagon secretion data for islets from donors without diabetes aged 15-39.**

Perifusion curves for HPAP UPenn (A) and HPAP Vanderbilt (B) data. Baseline glucagon secretion from UPenn (C) and Vanderbilt (D), area under the curve for amino acid-stimulated secretion from UPenn (E), low glucose and epinephrine-stimulated secretion from Vanderbilt (F), IBMX-stimulated secretion from UPenn (G) and Vanderbilt (H), and KCl-stimulated secretion from UPenn (I) and Vanderbilt (J). Islet glucagon content from UPenn (K) and Vanderbilt (L). AAM = amino acid mix (0.44 mM alanine, 0.19 mM arginine, 0.038 mM aspartate, 0.094 mM citrulline, 0.12 mM glutamate, 0.30 mM glycine, 0.077 mM histidine, 0.094 mM isoleucine, 0.16 mM leucine, 0.37 mM lysine, 0.05 mM methionine, 0.70 mM ornithine, 0.08 mM phenylalanine, 0.35 mM proline, 0.57 mM serine, 0.27 mM threonine, 0.073 mM tryptophan, and 0.20 mM valine, 2 mM glutamine), G = glucose, IBMX = 3-isobutyl-1-methylxanthine at 0.1 mM. KCl is at 30 mM for UPenn, 20 mM for Vanderbilt. Perifusion curves show mean ± SD. ns = not significant, $p \geq 0.05$.

### Supplement

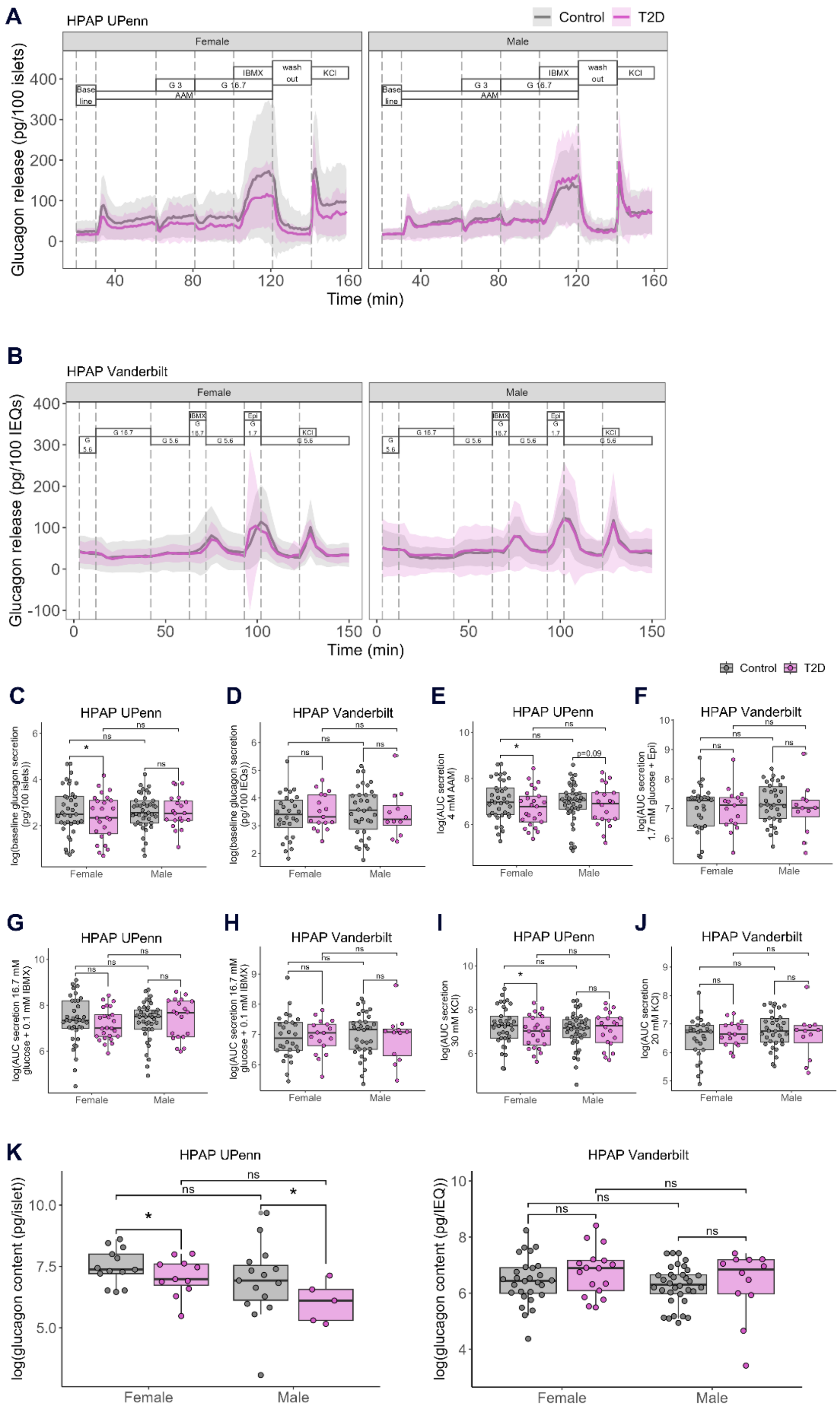
A
HPAP UPenn
Control
T2D
Female
Male
Base line
G 3
G 16.7
AAM
IBMX
wash out
KCl
Glucagon release (pg/100 islets)
Time (min)
B
HPAP Vanderbilt
G 5.6
G 16.7
IBMX
Epi
G 1.7
KCl
Glucagon release (pg/100 IEQs)
Time (min)
C
HPAP UPenn
log(baseline glucagon secretion (pg/100 islets))
D
HPAP Vanderbilt
log(baseline glucagon secretion (pg/100 IEQs))
E
HPAP UPenn
log(AUC secretion 4 mM AAM)
p=0.09
F
HPAP Vanderbilt
log(AUC secretion 1.7 mM glucose + Epi)
G
HPAP UPenn
log(AUC secretion 16.7 mM glucose + 0.1 mM IBMX)
H
HPAP Vanderbilt
log(AUC secretion 16.7 mM glucose + 0.1 mM IBMX)
I
HPAP UPenn
log(AUC secretion 30 mM KCl)
J
HPAP Vanderbilt
log(AUC secretion 20 mM KCl)
K
HPAP UPenn
log(glucagon content (pg/islet))
HPAP Vanderbilt
log(glucagon content (pg/IEQ))
ns
*
Female
Male

**Figure S18. HPAP dynamic glucagon secretion data for islets from donors without diabetes and donors with T2D of all ages.**

Perifusion curves for HPAP UPenn (A) and HPAP Vanderbilt (B) data. Baseline glucagon secretion from UPenn (C) and Vanderbilt (D), area under the curve for amino acid-stimulated secretion from UPenn (E), low glucose and epinephrine-stimulated secretion from Vanderbilt (F), IBMX-stimulated secretion from UPenn (G) and Vanderbilt (H), and KCl-stimulated secretion from UPenn (I) and Vanderbilt (J). Islet glucagon content from UPenn (K) and Vanderbilt (L). AAM = amino acid mix (0.44 mM alanine, 0.19 mM arginine, 0.038 mM aspartate, 0.094 mM citrulline, 0.12 mM glutamate, 0.30 mM glycine, 0.077 mM histidine, 0.094 mM isoleucine, 0.16 mM leucine, 0.37 mM lysine, 0.05 mM methionine, 0.70 mM ornithine, 0.08 mM phenylalanine, 0.35 mM proline, 0.57 mM serine, 0.27 mM threonine, 0.073 mM tryptophan, and 0.20 mM valine, 2 mM glutamine), G = glucose, IBMX = 3-isobutyl-1-methylxanthine at 0.1 mM. KCl is at 30 mM for UPenn, 20 mM for Vanderbilt. * indicates $p<0.05$, ns = not significant, $p \geq 0.05$.

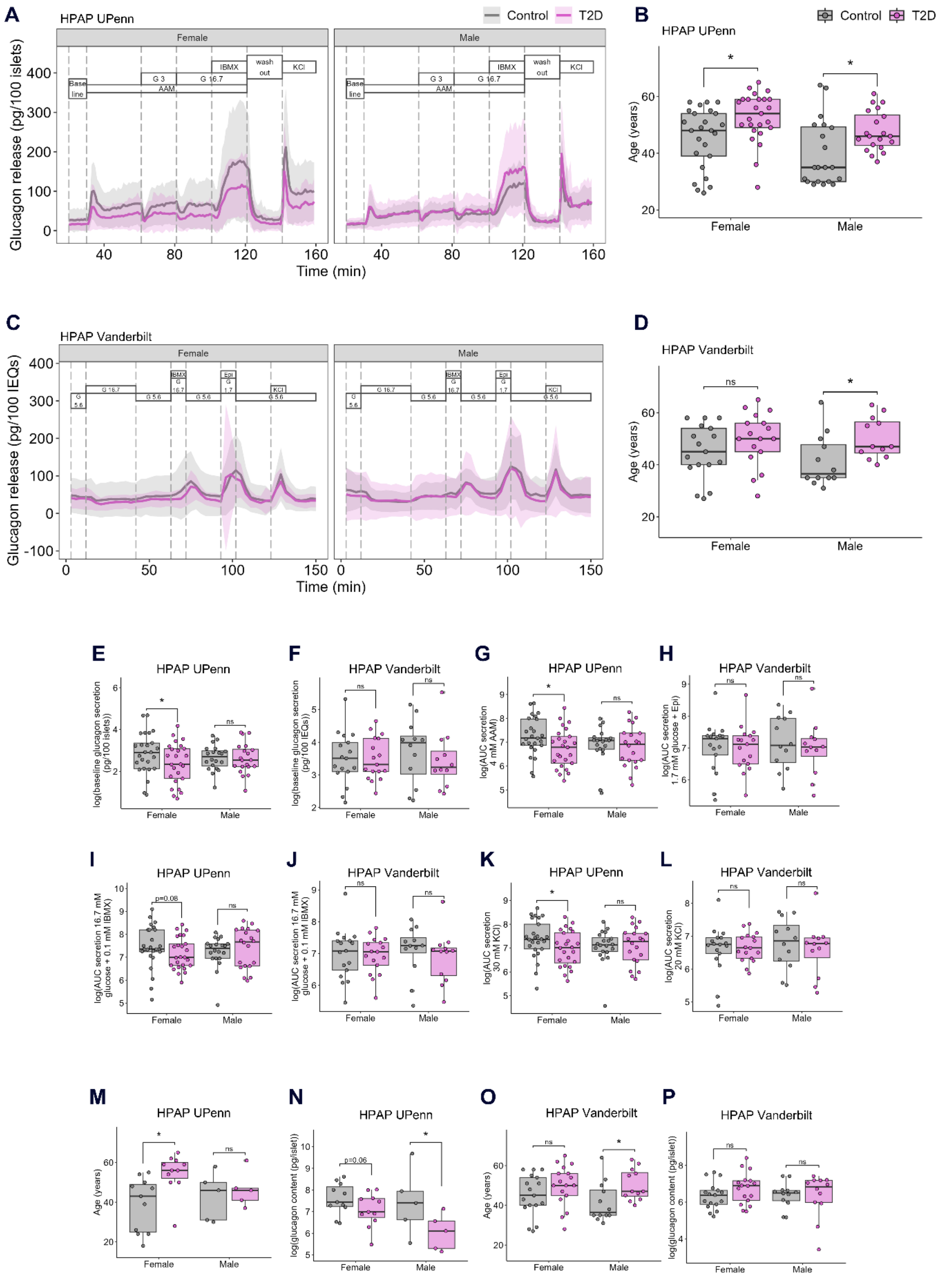
A
HPAP UPenn
Control
T2D
Female
Male
Base line
G 3
G 16.7
AAM
IBMX
wash out
KCl
Glucagon release (pg/100 islets)
Time (min)
B
HPAP UPenn
Age (years)
Female
Male
C
HPAP Vanderbilt
G 5.6
G 16.7
Epi
Glucagon release (pg/100 IEQs)
Time (min)
D
HPAP Vanderbilt
Age (years)
ns
E
HPAP UPenn
log(baseline glucagon secretion (pg/100 islets))
F
HPAP Vanderbilt
log(baseline glucagon secretion (pg/100 IEQs))
G
HPAP UPenn
log(AUC secretion 4 mM AAM)
H
HPAP Vanderbilt
log(AUC secretion 1.7 mM glucose + Epi)
I
HPAP UPenn
log(AUC secretion 16.7 mM glucose + 0.1 mM IBMX)
p=0.08
J
HPAP Vanderbilt
log(AUC secretion 16.7 mM glucose + 0.1 mM IBMX)
K
HPAP UPenn
log(AUC secretion 30 mM KCl)
L
HPAP Vanderbilt
log(AUC secretion 20 mM KCl)
M
HPAP UPenn
Age (years)
N
HPAP UPenn
log(glucagon content (pg/islet))
p=0.06
O
HPAP Vanderbilt
Age (years)
P
HPAP Vanderbilt
log(glucagon content (pg/islet))

200

**Figure S19. HPAP dynamic glucagon secretion data in age-matched data.**
Donors who lived with T2D were age-matched as closely as possible with donors without T2D within each sex. Perifusion curves (A) and donor ages (B) for age-matched HPAP UPenn data. Perifusion curves (C) and donors ages (D) for age-matched HPAP Vanderbilt data. Baseline glucagon secretion from UPenn (E) and Vanderbilt (F), area under the curve for amino acid-stimulated secretion from UPenn (G), low glucose and epinephrine-stimulated secretion from Vanderbilt (C), IBMX-stimulated secretion from UPenn (I) and Vanderbilt (J), and KCl-stimulated secretion from UPenn (K) and Vanderbilt (L) in age-matched data. Donor ages (M) and islet glucagon content (N) for age-matched UPenn data. Donor ages (O) and islet glucagon content (P) for age-matched Vanderbilt data. AAM = amino acid mix (0.44 mM alanine, 0.19 mM arginine, 0.038 mM aspartate, 0.094 mM citrulline, 0.12 mM glutamate, 0.30 mM glycine, 0.077 mM histidine, 0.094 mM isoleucine, 0.16 mM leucine, 0.37 mM lysine, 0.05 mM methionine, 0.70 mM ornithine, 0.08 mM phenylalanine, 0.35 mM proline, 0.57 mM serine, 0.27 mM threonine, 0.073 mM tryptophan, and 0.20 mM valine, 2 mM glutamine), G = glucose, IBMX = 3-isobutyl-1-methylxanthine at 0.1 mM. KCl is at 30 mM for UPenn, 20 mM for Vanderbilt. * indicates $p<0.05$, ns = not significant, $p\geq0.05$.

**Supplemental Table Legends**

**Table S1.** Full differential expression results for HPAP and Humanislets.com transcriptomics comparing female and male donors without diabetes aged 15-39.

**Table S2.** Full GSEA results for HPAP and Humanislets.com transcriptomics comparing female and male donors without diabetes aged 15-39.

**Table S3.** Full differential expression results for Humanislets.com proteomics comparing female and male donors without diabetes aged 15-39.

**Table S4.** Full GSEA results for Humanislets.com proteomics comparing female and male donors without diabetes aged 15-39.

**Table S5.** Full differential expression results for HPAP and Humanislets.com transcriptomics comparing female non-diabetic and T2D donors.

**Table S6.** Full differential expression results for HPAP and Humanislets.com transcriptomics comparing male non-diabetic and T2D donors.

**Table S7.** Full GSEA results for HPAP and Humanislets.com transcriptomics analyses comparing female non-diabetic and T2D donors.

**Table S8.** Full GSEA results for HPAP and Humanislets.com transcriptomics comparing male non-diabetic and T2D donors.

**Table S9.** Full differential expression for Humanislets.com proteomics comparing female non-diabetic and T2D donors.

**Table S10.** Full differential expression for Humanislets.com proteomics comparing male non-diabetic and T2D donors.

**Table S11.** Full GSEA results for Humanislets.com proteomics comparing female non-diabetic and T2D donors.

**Table S12.** Full GSEA results for Humanislets.com proteomics comparing male non-diabetic and T2D donors.
